# Ablation of Proliferating Osteoblast Lineage Cells After Fracture Leads to Atrophic Nonunion in a Mouse Model

**DOI:** 10.1101/2020.10.06.327288

**Authors:** Katherine R. Hixon, Jennifer A. McKenzie, David A.W. Sykes, Susumu Yoneda, Austin Hensley, Evan G. Buettmann, Hongjun Zheng, Dimitrios Skouteris, Audrey McAlinden, Anna N. Miller, Matthew J. Silva

## Abstract

Nonunion is defined as the permanent failure of a fractured bone to heal, often necessitating surgical intervention. Atrophic nonunions are a subtype that are particularly difficult to treat. Animal models of atrophic nonunion are available; however, these require surgical or radiation-induced trauma to disrupt periosteal healing. These methods are invasive and not representative of many clinical nonunions where osseous regeneration has been arrested by a “failure of biology”. We hypothesized that arresting osteoblast cell proliferation after fracture would lead to atrophic nonunion in mice. Using mice that express a thymidine kinase (tk) “suicide gene” driven by the 3.6Col1a1 promoter (Col1-tk), proliferating osteoblast lineage cells can be ablated upon exposure to the nucleoside analog ganciclovir (GCV). Wild-type (WT; control) and Col1-tk littermates were subjected to a full femur fracture and intramedullary fixation at 12 weeks age. We confirmed abundant tk+ cells in fracture callus of Col-tk mice dosed with water or GCV, specifically many osteoblasts, osteocytes, and chondrocytes at the cartilage-bone interface. Histologically, we observed altered callus composition in Col1-tk mice at 2 and 3 weeks post fracture, with significantly less bone and more fibrous tissue. Col1-tk mice, monitored for 12 weeks with *in vivo* radiographs and microCT scans, had delayed bone bridging and reduced callus size. Following sacrifice, *ex vivo* microCT and histology demonstrated failed union with residual bone fragments and fibrous tissue in Col1-tk mice. Biomechanical testing demonstrated a failure to recover torsional strength in Col1-tk mice, in contrast to WT. Our data indicates that suppression of proliferating osteoblast-lineage cells for at least 2 weeks after fracture blunts the formation and remodeling of a mineralized callus leading to a functional nonunion. We propose this as a new murine model of atrophic nonunion.

## Introduction

Nonunion is defined as the permanent failure of a fractured bone to heal, where surgical intervention is often required to achieve healing.^(1)^ The reported clinical rate of nonunion is 5%, with an estimated 500,000 fractures resulting in nonunion in the United States each year.^(2–4)^ Due to the necessary advanced care, treatment can cost upwards of $90,000 per individual.^(5)^ Nonunions are caused by a variety of factors such as infection, avascularity, or lack of stability, and are broadly categorized as hypertrophic or atrophic, each of which require different treatment options.^(1,6)^ Of these, atrophic nonunion (characterized by absent or minimal callus formation) is the least understood and the most difficult to treat.^(6,7)^ Available animal models for atrophic nonunion involve invasive methods such as local periosteal stripping, bone marrow removal, devascularization, or the creation of a critical-sized defect.^(6,8)^ Such invasive methods are not representative of many clinical nonunions which are due to the disturbance of biological pathways.^(1,9,10)^ Thus, there is an unmet need for a clinically relevant “failure of biology” atrophic nonunion animal model in which therapeutic interventions could be tested.

Proliferation of periosteal cells occurs during fracture healing, providing a source of callus osteoblasts and chondrocytes.^(6,11–13)^ Previous work in rodents has shown that cell proliferation in periosteal callus is elevated as early as 2 days and remains elevated through 14 days after fracture, as shown by expression of proliferating cell nuclear antigen (PCNA).^(14–16)^ These observations suggest that the first 2 weeks are a critical period for cell proliferation during rodent fracture healing. They further suggest that impaired cell proliferation during this period will lead to blunted callus formation, which may in turn result in atrophic nonunion. To our knowledge, it has not been shown that proliferation of periosteal progenitors in the early post-fracture period is required for successful healing.

While the progression of proliferation during fracture healing has been described, the molecular identity of these proliferating cells remains unclear. Recent reports of periosteal progenitors that contribute to fracture callus have used lineage tracing to identify a number of non-unique genes that mark this population.^(17)^ Earlier work demonstrated high GFP reporter expression in periosteal cells of 3.6Col1a1-GFP mice and showed that 3.6Col1a1 marks cells of the osteoblast lineage, from pre-osteoblasts to mature osteoblasts.^(18)^ An additional study further broadens this targeted population by demonstrating that in addition to osteoblast lineage cells, the 3.6 kb DNA fragment of the Col1a1 promoter also directs the expression of transgenes in the osteoclast lineage.^(19)^ We recently reported that proliferation of the 3.6Col1a1 cell population contributes to periosteal bone formation after non-injurious mechanical loading^(20)^, leading us to hypothesize that this population may also be critical to fracture healing, which is a largely periosteal-driven process.^(21)^ Jilka et al. ^(22)^ developed 3.6Col1a1-tk (Col1-tk) mice in which proliferating osteoblast lineage cells can be ablated through exposure to the nucleoside analog ganciclovir (GCV). In the presence of GCV, replicating cells expressing a thymidine kinase (tk) “suicide gene” convert GCV to a toxic nucleotide which is incorporated into the DNA resulting in targeted cell death.^(22)^ This model provides a unique tool to test the requirement of proliferation of a defined population of periosteal cells to fracture healing.

The central hypothesis of this study is that proliferation of periosteal osteoblast-lineage cells is required for fracture healing. We created midshaft femur fractures in young-adult Col1-tk mice, and treated them with GCV for 2 or 4 weeks to ablate proliferating osteoblast-lineage cells, followed by withdrawal. Wildtype (WT) littermate control mice were treated identically. Healing was assessed by *in vivo* serial radiography and microCT, followed by terminal assessment at 2-, 3- and 12-weeks post fracture using histology, microCT, and mechanical testing. Our findings show that suppression of proliferating osteoblast-lineage cells for either 2 or 4 weeks after fracture blunts the formation and remodeling of mineralized callus and leads to a non-functional fibrous nonunion. We propose this as a novel “failure of biology” murine model of atrophic nonunion.

## Materials and Methods

### Mouse Lines

A total of 128 male and female mice at 12 weeks of age were used in one of three experimental groups (**Figure 1A**). All experimental procedures were approved by the Institutional Animal Care and Use Committee (IACUC) at Washington University in St. Louis in accordance with the Animal Welfare Act and PHS Policy on Humane Care and Use of Laboratory Animals. Transgenic 3.6Col1A1-tk (Col1-tk) mice (provided by Drs. Robert Jilka and Charles O’Brien) were used to target replicating osteoblast progenitors.^(22)^ Specifically, these mice were developed to express the herpes simplex virus thymidine kinase (HSV-tk, or ‘tk’ for short) gene, driven by the 3.6 kb rat Col1A1 promoter which is active in osteoblast lineage cells. In the presence of the nucleoside analog ganciclovir (GCV), replicating osteoblast progenitors expressing thymidine kinase (tk) convert GCV to a toxic version of the nucleotide which is then incorporated into the DNA. Following integration, the DNA strands break, resulting in cell apoptosis (**Figure 1B**).^(22)^ To generate Col1-tk mice, male C57BL6/J (The Jackson Laboratory, #000664) mice were bred to female mice heterozygous for the tk transgene (tk-positive). This resulted in both heterozygous tk-positive (Col1-tk) and tk-negative (wildtype, WT) mice. Note that only one allelic copy of the tk transgene is necessary, and male Col1-tk mice are sterile. Genotyping was completed by Transnetyx using toe biopsies from the mice for real-time PCR (probe: puro). Subsequent breeding was completed using littermates (Col1-tk females and tk-negative males). All mice were group-housed up to five mice per cage under standard 12-hour light/dark cycle and given full access to food and water. Breeders were given high-fat chow and after weaning all mice were given normal chow. For the 2 week analysis timepoint only, two control groups were examined (WT dosed with GCV; and Col1-tk dosed with vehicle (water)), both of which should have normal osteoblast proliferation (**Figure 1B**). We chose WT dosed with GCV as the control group for all subsequent studies (3- and 12-week outcomes). These mice were dosed with ganciclovir (GCV, 8 mg/kg i.p., McKesson, San Francisco, CA) twice daily (**Figure 1A**) starting at the day of fracture (12 weeks of age). Mice were euthanized by CO2 asphyxiation at designated endpoints from 2 to 12 weeks after fracture.

**Figure 1.**
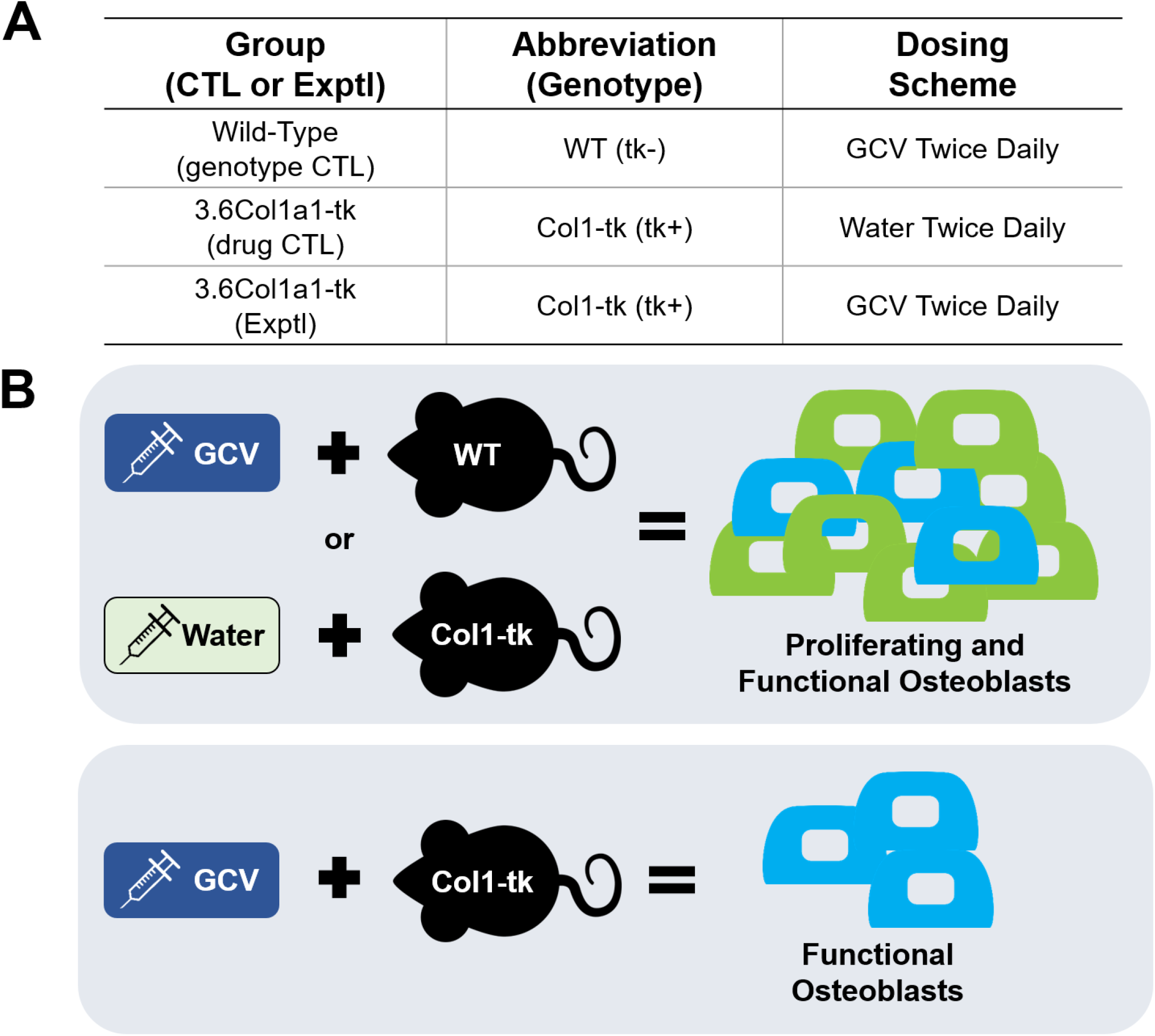
Experimental overview. Unilateral femur fractures were created in mice at 12 weeks age (N=128). (A) Both Col1-tk (tk-positive) and WT (tk-negative) mice were dosed with GCV, twice daily. Col1-tk (tk-positive) mice were also dosed with vehicle (water), twice daily. (B) In this experimental model, WT mice dosed with GCV and Col1-tk mice dosed with water have an increase in proliferating osteoblasts in addition to the resident functional osteoblasts. In contrast, Col1-tk mice only have resident functional osteoblasts present due to the ablation of proliferating osteoblasts with GCV administration.

### Fracture Healing Study

In healthy mice, complete bridging of the fracture site with bony callus occurs 3-4 weeks after fracture.^(23,24)^ In humans, fractured long bones typically heal within 2-3 months and nonunions are clinically diagnosed after 6-9 months of nonhealing, i.e., 3-times the normal healing time.^(23,25)^ Thus, it has been proposed that a conservative assessment of nonunion in mice should be based on evaluation at 12 weeks post fracture.^(23,26)^ In a subset of mice (n=19), WT mice were dosed with GCV and Col1-tk mice were dosed with either GCV or vehicle (water) for 2 weeks after fracture. Immunohistochemistry on decalcified, paraffin-embedded sections was used to visualize expression of tk in fracture callus, along with collagen 2, collagen 3, endomucin, and tartrate-resistant acid phosphatase (TRAP), as detailed below. For the main aim of the study, mice were dosed with GCV following fracture to target both the proliferative phase (2 week dosing) as well as the interval during which normal fracture healing occurs (4 week dosing). Following dosing for 2 or 4 weeks, GCV was then withdrawn and weekly monitoring of fracture healing continued through sacrifice (3 or 12 weeks) to evaluate healing. Following euthanasia, animals were randomly assigned to either histological or biomechanical evaluation.

### Fracture Model

A previously established protocol was used to create full fracture in the right femur of 12-week-old mice.^(27)^ Briefly, a transverse force was applied across the right femoral mid-diaphysis using a custom 3-pt bending setup (DynaMight 8841; Instron) creating a fracture. Following injury, the femur was stabilized using a 24- gauge metal pin (MicroGroup) and the wound closed using 3-0 nylon sutures (Ethicon). Fixation was confirmed by radiograph (Faxitron Ultrafocus 100; Faxitron Bioptics) immediately following surgery. This model mimics classical fracture injury and healing ^(28,29)^, with both intramembranous bone formation and endochondral healing around the periphery and within the fracture gap, respectively.^(24,30)^ All mice were given buprenorphine SR (1 mg/kg, s.c.) for analgesia 1 hour before surgery.

### Radiographic Evaluation

Lateral radiographs were taken weekly following full fracture until euthanasia (3X magnification; Faxitron UltraFocus100; n=12-18/group). All radiographs were blindly scored for the degree of healing using a modified Goldberg score (i.e. 0 = no bridging, 1 = one side bridged, 2 = complete bridging).^(31)^ Mice were excluded from the study if there was a loss of fixation during healing (total 18 mice).

### Histological Analysis

Femurs designated for paraffin processing (n=4-8/group/timepoint) were dissected and fixed immediately in 10% neutral buffered formalin for 24 hours, followed by decalcification in 14% EDTA (pH 7.0) for 2 weeks. Standard paraffin processing was used such that fractured femurs were cut as longitudinal sections at a 5 μm thickness. Slides were stained with H&E, Movat’s pentachrome and picrosirius red/alcian blue at various timepoints post fracture to get a general overview of the different tissues involved in fracture healing. These stains can differentiate the mineralized callus, cartilage maturation, and fibrous tissues with different colors. Movat’s pentachome stains the mineralized bone (yellow), cartilage ground substance (green), fibrin-fibrinoid, and muscle (red). Alcian blue stains the glycosaminoglycans and ground substance mucins (cartilage and chondrocytes) (blue). The picrosirius red stains the collagen (red). Immunohistochemistry (IHC) was used to localize HSV-tk expression, along with collagen 2 (col2, cartilage matrix), collagen 3 (col3, fibrous tissue and new vessels), endomucin (vessels), and tartrate-resistant acid phosphatase (TRAP, osteoclasts).

Briefly, the paraffin slides were deparaffinized in xylene and rehydrated in graded ethanols. Incubation with 3% H_2_0_2_ (5 min) blocked endogenous peroxidases. Proteinase K was used for antigen retrieval (col3 10 min at 37 C, col2 35 min at 37C, endomucin 5 min 23 C) prior to blocking step. HSV-tk and col3 endogenous epitopes were blocked with 10% goat serum (abcam – ab7481) in PBS at room temp (1 hour). The sections were then incubated with rabbit polyclonal anti-HSV tk (1:100; gift from William Summers, Yale) or rabbit polyclonal to col3 (1:500, abcam ab7778) overnight at 4C. Both HSV-tk and col3 used a secondary polymer-HRP anti-rabbit antibody incubation for 1 hr (1:200 Dako P0448). For col2 and endomucin staining the manufacturer’s instructions were followed (Vectastain Elite ABC kit, Rat IgG PK-6104) with primary antibodies for col2 (1:100; gift from Linda Sandell, WashU) or endomucin (1:400, eBioV.7C7 #14-5851-85). All IHC samples were incubated with DAB chromagen (brown; Vector DAB, SK-4105, 30-60 second). Sections were counterstained with Modified Mayer’s hematoxylin (Vector Hematoxylin QS, H-3404) and imaged at 20x on a Nanozoomer slide scanner (Hamamatsu Photonics). Isotype control antibodies were used as a negative controls (col2/Endomucin: eBioscience, Clone eBR2a; col3: Cell signaling, 2729S).

Endomucin analysis was done in four 20x ROI’s (one from each quadrant of the callus section) from which both woven bone and vessel area were quantified. The presence of osteoclasts was assessed using TRAP staining on paraffin sections. Slides were manually evaluated to assess osteoclasts on the peripheral surface of the callus (osteoclast length/callus length (%)) and the percent of the callus surface occupied by cartilage, fibrous tissue, and bone was quantified. Note that for the 3 week animals, the periphery of the entire callus (top and bottom) was analyzed for both WT and Col1-tk mice and varied in size, ranging from 4.8 to 14.8 mm. In 12 week animals, a 2 mm region spanning the original fracture site was analyzed for all mice due to the WT mice having healed by this timepoint. Picrosirius Red/Alcian Blue images were blinded and manually assessed quantitatively for callus composition where the entire callus was analyzed for 2 and 3 week data and a 4 mm ROI centered at the fracture site was used for 12 week data. For the 2 week timepoint the callus was separated into three tissue types based on staining and cell morphology: bone (dark red), cartilage (blue) and fibrous tissue (pale red)/other. At the 3- and 12-week timepoints the callus was assessed for bone, cartilage, fibrous/other, and periosteum (added to account for the thick periosteal tissue evident especially on the Col1-tk samples). Marrow area was not counted separately but included with ‘bone’.

To assess proliferation following GCV drug withdrawal, 5-Ethynyl-2’-deoxyuridine (EdU, 0.2 mg/mL in 5% sucrose) was added daily to the drinking water of a subset of mice. For 2 weeks after fracture mice were treated with GCV but not EdU. During the next (third) week, mice received EdU but not GCV. After this third week, the mice were sacrificed to analyze the cumulative proliferative response (EdU staining), i.e., during the week after removal of the anti-proliferative conditions. Femurs from mice given EdU designated for frozen processing following fracture (n=7/group) were dissected and fixed immediately in paraformaldehyde for 48 hours, followed by decalcification in 14% EDTA (pH 7.0) for 2 weeks. Samples were rinsed in PBS, infiltrated in 30% sucrose, and embedded in optimal cutting temperature compound (OCT compound, Tissue-Tek; VWR). A Cryojane Tape Transfer System (Leica) was used to create frozen longitudinal sections of the decalcified bone (Leica CM 1950 Manual Cryostat) at 5 μm thickness. Following sectioning, Click-iT EdU Alexa Fluor 647 Imaging Kit from ThermoFisher (C10340) was used to stain the sections for EdU and 4′,6-diamidino-2-phenylindole (DAPI; #D9542; 1:1000 dilution; Sigma-Aldrich, St. Louis, MO, USA). Briefly, frozen sections were thawed, washed with PBS, and permeabilized in 0.5% Triton X-100 (in PBS) for 20 min. The samples were then rinsed with PBS and incubated with 100 μL of the reaction cocktail (1x Click-iT reaction buffer, CuSO4, 1x buffer additive, and Alexa Flour azide) for 30 min. Following additional rinsing in PBS, the slides were counterstained with DAPI. The whole femur was imaged using the Zeiss Axio Scan.Z1 slide scanner (20x objective). A 4 mm ROI for both the top and bottom callus (excluding cortical bone and marrow) were exported and the area (px^2^) of both DAPI (blue) and EdU+ (pink) stained cells were assessed using ImageJ.

### Micro-Computed Tomography

*In vivo* scans were performed at 2, 4, 8, and 12 weeks, or 4, 6, 10, and 12 weeks for 2 and 4 weeks of GCV dosing, respectively (n=7-9/group). For each scan, the animal was anesthetized (1-3% isoflurane gas) and both femurs were scanned simultaneously using microCT (VivaCT 40, Scanco Medical AG, Switzerland; 15 μm voxel size, 70 kV, 114 μA, and 300 ms integration time). This resulted in an estimated local radiation exposure of 42 cGy/scan; no impairments in callus healing in WT mice were noted.^(32)^ Due to the artifact from the metal pin, threshold- or density-based measurements using Scanco software could not be performed. The proximal and distal ends of the callus were visualized, and a measurement of 3D distance between the center of the two sections was calculated (mm). Contour lines were drawn around the outer edge of the callus and the callus volume was output as total tissue volume (TV, mm^3^). For *ex vivo* scans, femurs were dissected from the surrounding tissue and fracture fixation pins were removed (n=15-18/group). Femurs were scanned using microCT (VivaCT 40, Scanco Medical AG, Switzerland; 10.5 μm voxel size, 55 kV, 145 μA, 300 ms integration time). Analysis regions of interest (ROIs) differed for *ex vivo* 3 and 12 week femurs due to changes in callus length over time. For femurs dissected at 3 weeks (n=10-12/group), a 600 slice (6.3 mm length) ROI was centered at the midpoint of the fracture line and a threshold of 200 per mille was used. The ROI length was selected to include the majority of callus region of all samples. Each callus was contoured around its periphery, and the ROI included the original cortical bone plus the callus. Outcomes were total bone volume (BV, mm^3^), tissue volume (TV, mm^3^), bone volume per total volume (BV/TV, mm^3^/mm^3^), and tissue mineral density (TMD, mg HA/cm^3^). At 12 weeks post fracture (n=14-17/group), a region of interest (ROI) of 200 total slices (2.1 mm) was identified, centered at the point of fracture, and the threshold was set to 350 per mille. This smaller ROI length reflected the shorter extent of mineralized callus at this timepoint. Due to remodeling of the callus, fracture lines were not visible at 12 weeks, thus the point of fracture was inferred from radiographic images at 2 weeks post fracture (femoral head to fracture midpoint distance). Contours around the periosteal bone surface were drawn and BV, TV, volumetric bone mineral density (vBMD, mgHA/cm^3^), and TMD were determined. Following *ex vivo* scanning, the femurs were either decalcified and processed for histology, or prepared for mechanical testing.

### Limb Bud Assay

High-density micromass cultures from murine limb buds were utilized to study the effect of GCV on chondrogenesis and osteogenesis, based on a previously established protocol.^(33,34)^ The hind limb buds of WT and Col1-tk embryos (E11-E12) were harvested and rinsed in HBSS at room temperature (n=4-8/group). Following two PBS washes, the cells were centrifuged (6,000 rpm, 2 min), digested (1X trypsin), and incubated at 37 °C for 2-3 min. The limb buds were then dissociated by gentle pipetting and quenched with 100 μl DMEM. The cell suspension was filtered (40 μm nitex) and centrifuged at 6000 rpm for 2 min. The cells were then resuspended in DMEM, counted, and reconstituted at 1 − 2 × 10^7^ cells/ml. 20 μL of cells (2 − 4 × 10^5^ cells) from each sample were seeded in the center of two wells of a 12-well plate and allowed to attach for 10 – 20 min. Note that this range of cells may account for slight variability in micromass size between samples. 1-2 mL DMEM was added to each well and media was changed every other day for 6 days. Immediately following initial seeding, one well for each sample was dosed with 10 μg/mL of GCV, based on a previous study and pilot work.^(35)^ At day 6, half of the samples were rinsed with PBS, fixed with 3% glacial acetic acid in 100% ethanol, rinsed again with PBS, and stained with Alcian Blue dye overnight. All remaining samples were given media supplemented with 10 mM β-glycerophosphate and 50 μg/mL ascorbic acid to promote osteogenic differentiation (prepared fresh; changed every other day) through day 16. At this time the samples were rinsed with PBS and stained with Alizarin Red for 30 minutes. Brightfield images at day 16 displayed cell morphology and mineral formation. Additional images of the wells were also taken following Alizarin Red staining.

### Biomechanical Testing

Bilateral femurs were dissected at 12 weeks post fracture and cleaned of all soft tissue (n=6-9/group). The ends of each femur were potted using polymethylmethacrylate (PMMA, Ortho-Jet, Land Dental) in 6mm diameter × 12mm length acrylic tubes. The bone was centered using a custom fixture, leaving approximately 4.2 mm of exposed bone (including the callus region) between potting tubes. All samples were wrapped in PBS soaked gauze to preserve hydration while the PMMA cured overnight. The following day, each sample was loaded into a custom-built torsion machine with a 25 in-oz load cell controlled with LabVIEW software (LabVIEW 2014, National Instrument, TX). The machine held one of the potted femur ends in a fixed position while rotating the other potted tube at 1 deg/sec until fracture. The maximum torque (Nmm), rotation at maximum torque (degrees), and stiffness (Nmm/degree) were calculated from the resulting torque-rotation graphs (Matlab).

### Statistics

Prior to experiments, study sample sizes were calculated based on a power analysis with α = 0.05 and β= 0.20 (https://www.stat.ubc.ca/~rollin/stats/ssize/n2.html). Estimates of sample variance and effect size were based on previous experimental data and biological importance.^(24,30)^ Target samples sizes for outcomes per experimental group were: MicroCT: n = 8, Histology: n = 7, Biomechanics: n = 10. Actual sample sizes are noted above and in Results. A chi-square test was used to assess fracture union based on the Goldberg scale for each timepoint. An unpaired t-test was used to compare WT and Col1-tk microCT data taken at 3 weeks after fracture. Picrosirius Red/Alcian Blue and TRAP histological quantification taken at 2, 3, and/or 12 weeks post fracture was also analyzed using an unpaired t-test. Two-way ANOVA with Sidak’s post hoc test was used to analyze Endomucin and Picrosirius Red/Alcian Blue at 3 and/or 12 weeks post fracture. Two-way ANOVA with Sidak’s post hoc test to correct for multiple comparisons was used for *in-vivo* microCT (repeated factor: *time*; between factor: *genotype*). *Ex-vivo* microCT and torsion testing data were also analyzed using two-way ANOVA with Sidak’s post hoc test (repeated factor: *side* [intact, fractured]; between factor: *genotype*). Significance was considered at p values < 0.05. All data analysis was performed using Prism (Version 8; GraphPad Software, La Jolla, CA, USA).

## Results

### Localization of tk+ Cells in Fracture Callus

To assess which cells were being targeted, HSV-tk, col2, col3, and endomucin expression was evaluated in fracture callus of Col1-tk mice dosed with GCV or water at 2 weeks post fracture. In Col1-tk mice dosed with vehicle (water), callus composition appeared normal and H&E stained callus sections displayed tissues from three general areas of the callus: 1) bone, 2) cartilage and 3) fibrous/other (**Figure 2A**). Tk+ expression was noted in cells within the woven bone and cartilage regions and in periosteal cells overlying the callus. Overall, cells positive for HSV-tk predominately resided within bone tissue (osteoblasts and osteocytes). HSV-tk+ cells were observed on the periphery of cartilage islands, but no HSV-tk+ cells were detected within the middle of these areas. Some HSV-tk+ cells were detected within fibrous tissue areas, but these cells did not consistently correlate with col3 positive areas. Comparatively, Col1-tk mice dosed with GCV for 2 weeks after fracture displayed a reduced fracture callus with minimal woven bone and cartilage (**Figure 2B**). Similar to control fractures, HSV-tk staining occurred in cells within and around the bone matrix. In cartilage areas that were beginning to mineralize, HSV-tk+ cells were found on the periphery in areas that had mineral staining (yellow on pentachrome, red on picosirius red). In some fibrous/other tissue areas there was evidence of HSV-tk+ cell staining; these cells did not stain positive for col2 or col3. WT mice given GCV displayed typical callus composition with both woven bone and cartilage, but no tk+ staining (data not shown).

**Figure 2.**
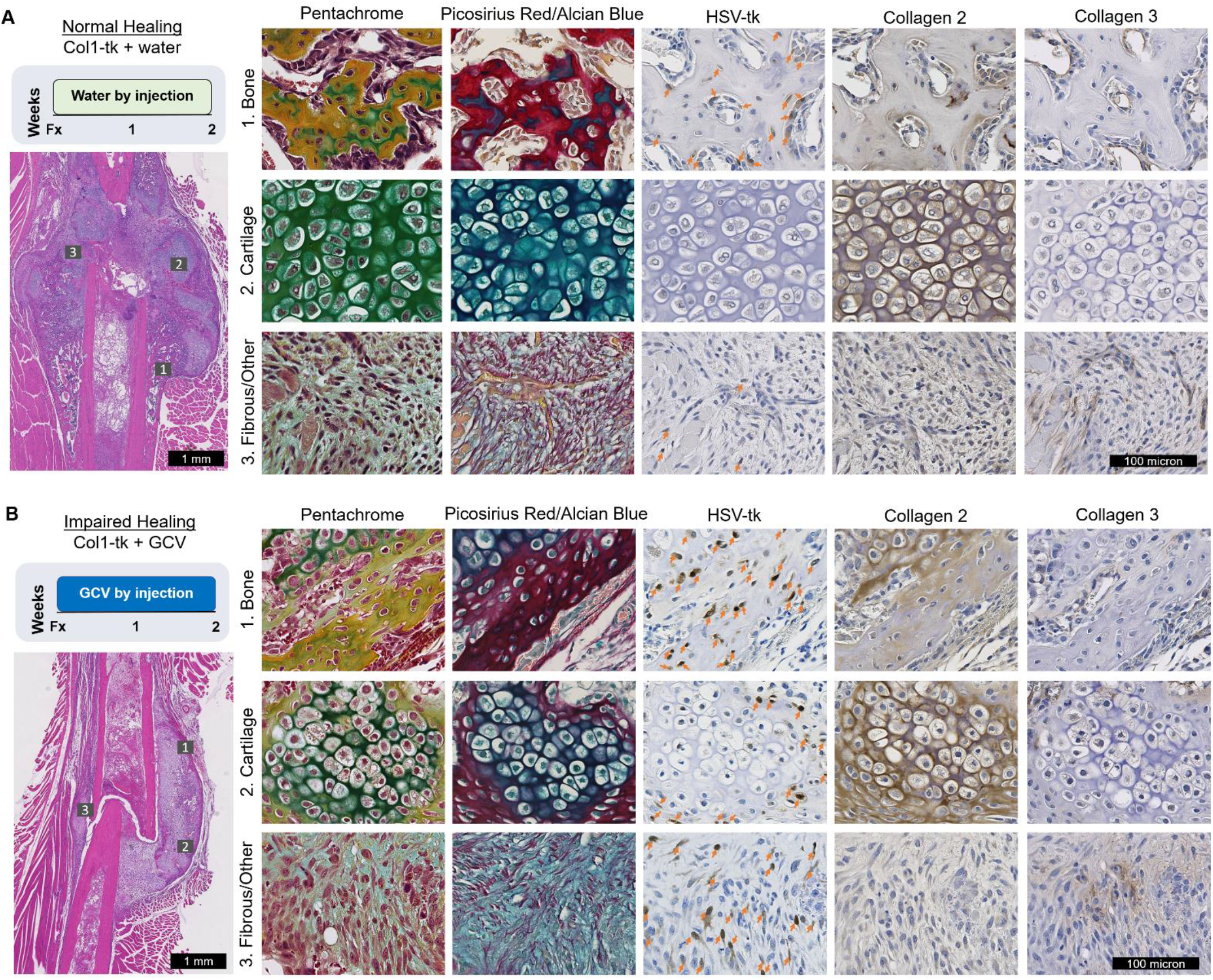
Histological characterization of Col1-tk positive cells within different areas of the fracture callus (n=4-8/experimental group/timepoint). (A) Normal fracture healing occurs in a Col1-tk+ mouse with vehicle (water) injections for two weeks after surgery. H&E stained callus sections at 2 weeks post fracture were used to identify tissues from three different areas of the callus: 1) bone, 2) cartilage and 3) fibrous/other. Histological staining was completed on serial sections for Movat’s pentachrome, picosirius red and alcien blue, HSV-tk, collagen 2 and collagen 3. Cells positive for HSV-tk (orange arrows) predominately resided within bone tissue (osteoblasts and osteocytes). No HSV-tk+ cells were detected within the middle of large cartilage areas. Some HSV-tk+ cells were detected within fibrous tissue areas, but these cells did not seem to consistently correlate with collagen 3 positive areas. (B) Healing was impaired with the addition of ganciclovir (GCV) drug for two weeks after fracture. Tissue distributions were different in the impaired healing samples, but most samples had some areas of bone, cartilage and fibrous/other to examine. Similar to control fractures, HSV-tk staining was most evident in cells within and around the bone matrix. In cartilage areas that were beginning to mineralize HSV-tk+ cells were found on the periphery, coincident with areas that had mineral staining (yellow on pentachrome, red on picosirius red). In some fibrous/other tissue areas there was evidence of HSV-tk+ cell staining. These cells did not stain positive for collagen 2 or 3 and were not located within distinct cartilage or bone areas of the callus.

### Bony Bridging from 1 to 3 Weeks Was Reduced for Col1-tk Mice

Early healing was evaluated following 2 weeks of GCV dosing and 1 week of withdrawal by weekly radiographs, which were blindly scored using a modified Goldberg score (**Figure 3A**). At 1 week after fracture, almost no bridging had occurred in either genotype. At 2 weeks, over 80% of the WT mice had at least one side bridged, compared to just over 50% of Col1-tk mice. After 3 weeks, there was some bridging in 100% of WT mice, and 83% had complete bridging; this was significantly different than Col1-tk mice, where there was some bridging in 77% of samples, but no samples were completely bridged, and 23% remained unbridged (χ^2^, p < 0.0001; **Figure 3B**). Representative radiographic images demonstrate good and poor healing outcomes in WT and Col1-tk mice, respectively (**Figure 3C**). Additionally, 3D reconstructions of *ex vivo* microCT scans show a robust callus formed at the fracture site of WT femurs while the Col1-tk femurs showed greatly reduced callus volume (TV) and bone volume (BV) (**Figure 3C**). Quantitative analysis of these scans (including callus and cortical bone) demonstrate an approximate 50% reduction in BV and TV from WT to Col1-tk (p < 0.001; **Figure 3D, E**). Measures of density, i.e., BV/TV and TMD, were greater in Col1-tk compared to WT femurs (p < 0.01), reflecting a larger relative contribution of the original (dense) cortical bone to the total bone within the ROI **(Figure 3F, G**). This is supported by observations that the BV, TV, BV/TV, and TMD values for Col1-tk calluses were closer to the average values for intact femurs (shaded dashed line **Figure 3D-G**). Finally, analysis of the intact femurs by microCT showed no differences between Col1-tk and WT bones, indicating no effect of targeted ablation on uninjured bones (**Supplementary Table 1**).

**Figure 3.**
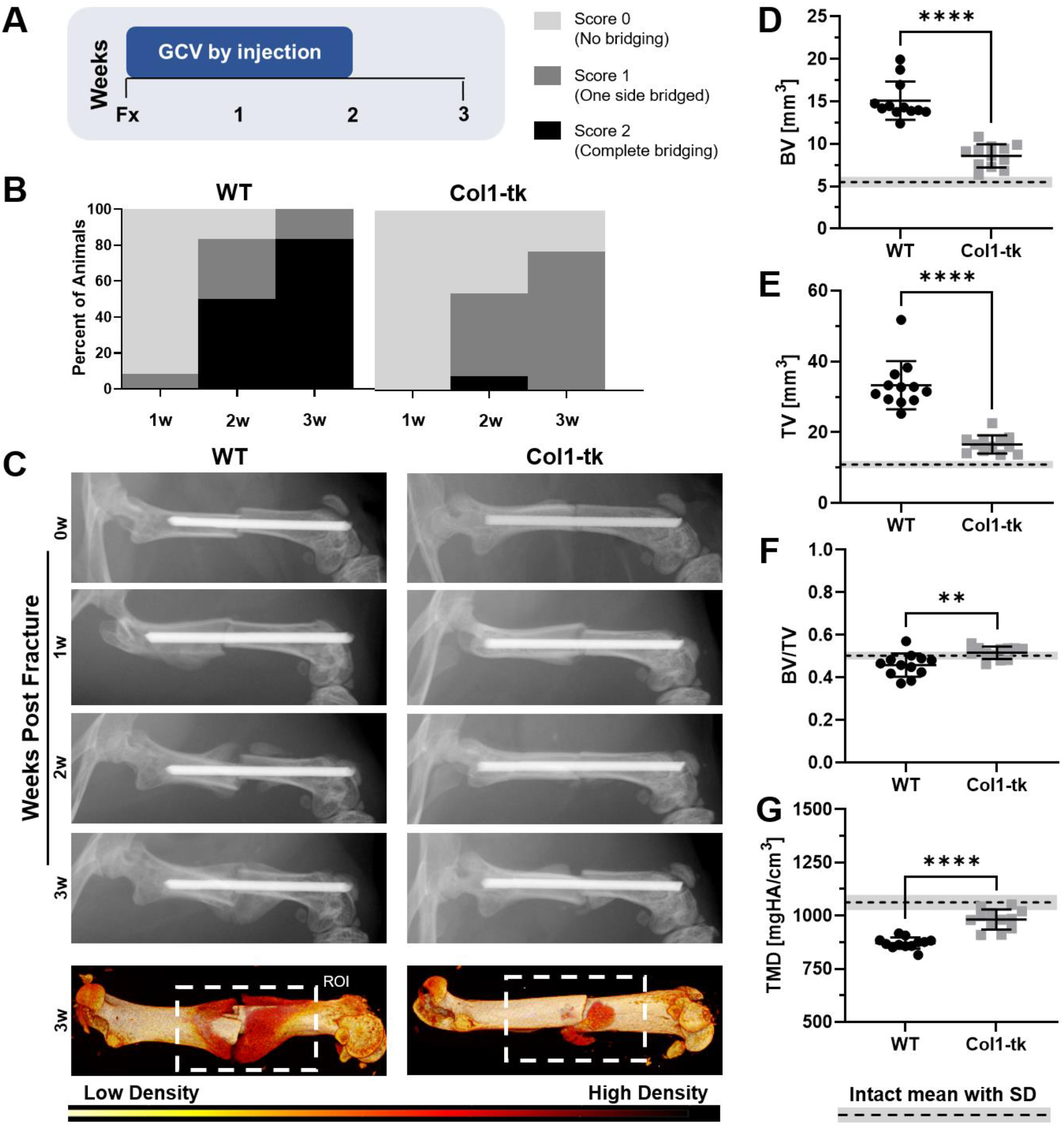
Following fracture, early (3 week) healing was evaluated by weekly X-rays as well as *ex vivo* microCT (n=12-18/group). (A) Mice were dosed with GCV for 2 weeks, followed by one week of drug withdrawal prior to sacrifice. (B) Bridging evaluated by a modified Goldberg scale demonstrated the progression of callus bridging in WT mice as compared to reduced or absent bridging in Col1-tk mice. (C) This lack of bridging and reduced callus formation in Col1-tk mice can also be visualized in both radiographic images and microCT 3D reconstructions (ROI = 6.3 mm region of interest for microCT analysis includes cortical bone and callus). MicroCT quantification showed significantly less (D) BV (bone volume) and (E) TV (tissue or callus volume) in Col1-tk vs. WT mice, but significantly higher (F) BV/TV and (G) TMD (tissue mineral density) (**p < 0.01, ***p < 0.001). Average values of BV, TV, BV/TV, and TMD for intact femurs are noted by a dashed line, where shading designates standard deviation.

### Callus Composition Was Altered in Col1-tk Mice at 2 and 3 Weeks

To further analyze healing, histological evaluation at 2 and 3 weeks after fracture was completed using Picrosirius Red/Alcian Blue staining (**Figure 4**). The femurs of the WT mice and the Col1-tk dosed with water appeared fully (or nearly fully) bridged by a large woven bone callus, with some cartilage present. There were no significant differences between the two control groups (WT dosed with GCV or Col1-tk dosed with water) at 2 weeks. In the group which arrested osteolineage cell proliferation, Col1-tk dosed with GCV had significantly less bone formation and significantly more fibrous tissue/other at the fracture site (**Figure 4A**). At the 3 week timepoint (1 week post GCV withdrawal), WT fracture sites contained significantly more bone and significantly less fibrous tissue than the Col1-tk mice (**Figure 4B**). Additional staining at 3 weeks showed that all tissue types were present in both WT and Colt-tk samples, and that the pattern of HSV-tk expression was similar to week 2 (**Supplementary Figure 1)**. HSV-tk+ staining was found within and around the bone tissue, on the mineralizing margins of the callus, and within some fibrous tissue areas of the Col1-tk samples. The HSV-tk+ cells did not appear to be positive for col3, TRAP, or endomucin.

**Figure 4.**
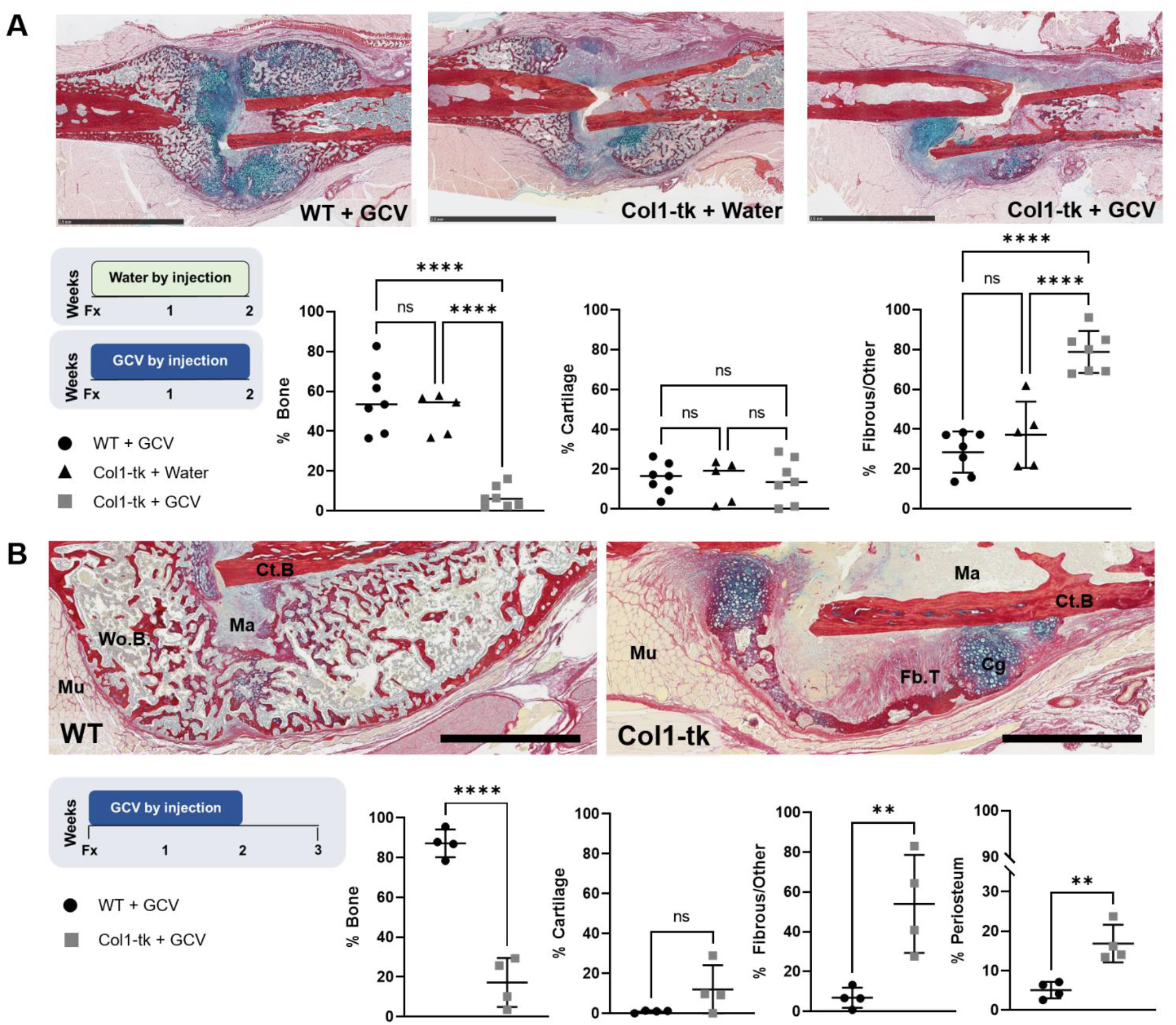
Healing at (A) 2 and (B) 3 weeks post fracture was qualitatively assessed using histological staining with Picrosirius Red/Alcian Blue (n=4-7/group). (A) At 2 weeks post fracture, sagittal sections from the fracture midpoint of WT+GCV and Col1-tk+water mice had cartilage (Cg; blue stain) and early woven bone (Wo.B; red stain) formation throughout the callus. The Col1-tk sections had a smaller callus composed of mostly cartilage (Cg; blue stain) and fibrous tissue (Fb.T/other), with little woven bone. Statistical analysis showed significantly less bone and significantly more Fb.T/other in the Col1-tk mice as compared to the controls (***p < 0.001). (B) Similar results were shown at 3 weeks post fracture, with WT mice now exhibiting complete bridging with the callus almost entirely composed of woven bone (Wo.B; red stain). The Col1-tk sections still had a smaller callus composed of primarily cartilage (Cg) and fibrous tissue (Fb.T/other), with little woven bone. Statistical analysis again showed significantly less bone and significantly more Fb.T/other in the Col1-tk mice as compared to the WT mouse (**p < 0.01, ***p < 0.001). Abbreviations: Cortical Bone = Ct.B,; Cartilage = Cg; Fibrous Tissue = Fb.T.; Marrow = Ma; Muscle = Mu; Woven Bone = Wo.B. Black scale bars denote 1 mm.

### Col1-tk Limb Bud Cells Treated with GCV Showed Slightly Reduced Chondrogenesis While Osteogenesis was Strongly Impaired

The *in vitro* limb bud assay focused at early (day 6) and later (day 16) timepoints to assess chondrogenesis and osteogenesis, respectively, in WT and Col1-tk mice (**Supplementary Figure 2**). When WT and Col1-tk cells are not dosed with GCV, there is no difference between micromass size and Alcian Blue staining. The area of the Alcian Blue stained region appeared slightly smaller in the Col1-tk group when comparing GCV treated cultures versus control untreated cultures, an effect not observed with the WT group. When the cells were cultured in an osteogenic induction media and stained with alizarin red on day 16, the three control groups demonstrate similar staining (WT + no GCV, WT + GCV, Co1-tk + no GCV). However, the Col1-tk + GCV group has reduced alizarin staining, as well as differences in cell morphology and mineral formation as observed under brightfield view.

### Proliferation was Robust in Fracture Callus

EdU/DAPI staining was used to characterize proliferation at 3 weeks (one week after GCV withdrawal) in the periosteal callus (**Supplementary Figure 3**). While the WT mice visually had more pink (EdU+) staining in some callus areas (**Supplementary Figure 3A’-B’**), the total quantified proliferating area was not significantly different from the Col1-tk mice (**Supplementary Figure 3C**). The WT mice had significantly more DAPI stained cells at the fracture site, which is consistent with its larger callus (**Supplementary Figure 3C**). When the ratio of EdU+ cells to DAPI+ cells is compared between WT and Col1-tk mice, there is no significant difference with both averaging approximately 35% proliferation of the total cells present in the fracture callus (**Supplementary Figure 3D**). Thus, in the week after GCV withdrawal, the rate of proliferation appears normal in the Col1-tk fracture callus.

### There was Reduced Vasculature and Osteoclasts in Col1-tk Calluses at 3 Weeks

TRAP and endomucin staining were performed to assess the presence of osteoclasts and vasculature, respectively, at 3 weeks (**Figure 5**). WT mice displayed an abundance of TRAP+ woven-bone lining osteoclasts along the callus surface (**Figure 5B, B’, B”**). While the Col1-tk mice also had osteoclasts lining some woven bone (**Figure 5C, 5C”**), this was not seen consistently throughout the callus (**Figure 5C’**). Osteoclast length per callus length was significantly greater in WT compared to Col1-tk mice (22% vs. 0.8%, p < 0.001; **Figure 5D**). Endomucin staining revealed a large amount of vasculature throughout the entire callus in the WT mice (**Figure 5E**). Comparatively, while the Col1-tk mice had some vasculature, a majority of the fracture site displayed little to no endomucin staining (**Figure 5F**). When analyzing four 20x fields from different quadrants of the callus, WT mice had significantly more woven bone and vasculature within the fracture callus than the Col1-tk mice (**Figure 5G**). Interestingly, when only the woven bone regions of the callus were quantified, Col1-tk mice still had significantly less vasculature, demonstrating that this difference is not solely due to a smaller fracture callus or differences in callus composition (**Figure 5H**).

**Figure 5.**
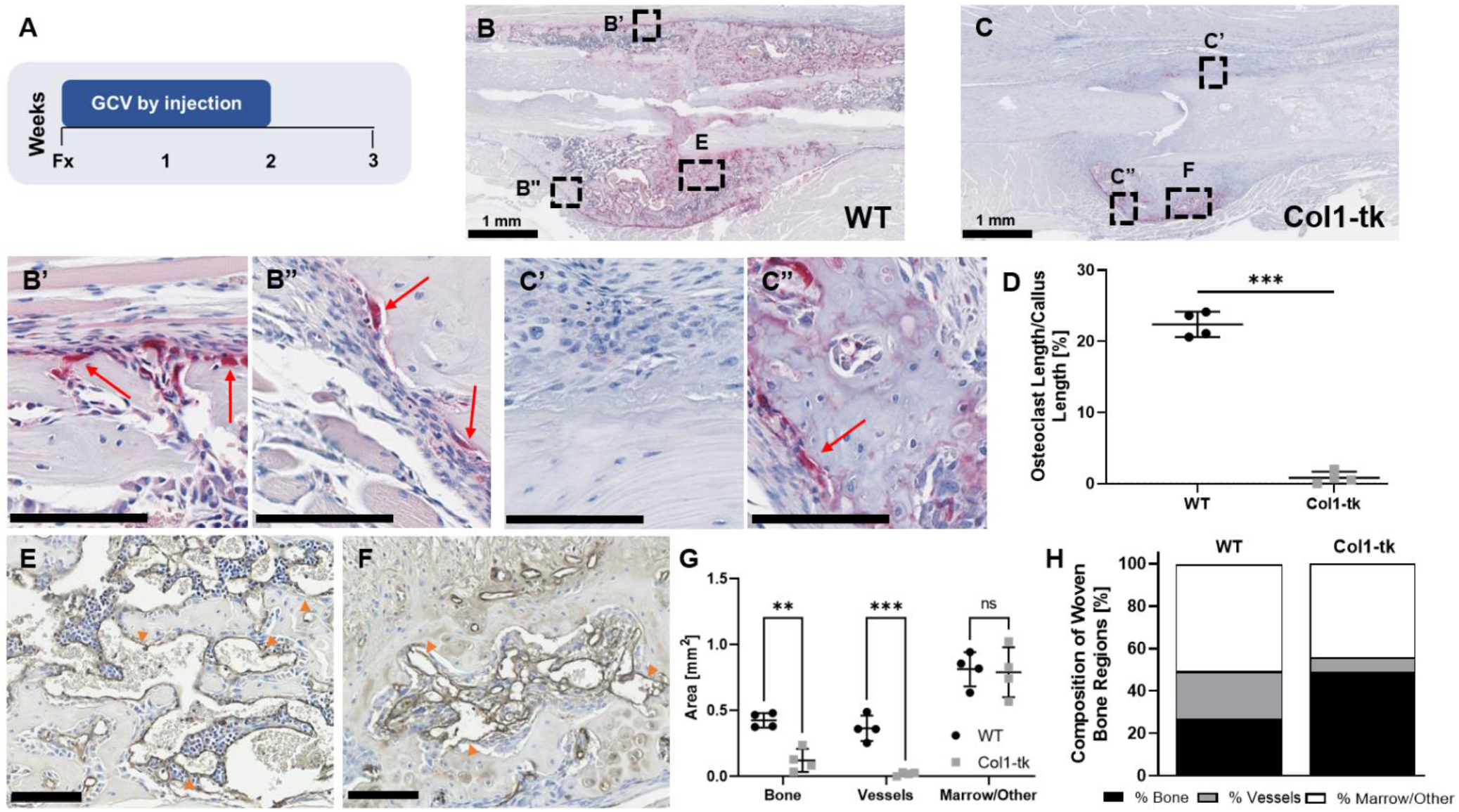
(A) Osteoclast (TRAP+) activity and the presence of vessels was evaluated in WT and Col1-tk mice at 3 weeks post fracture (n=4/group). (B) WT mice displayed many (red) TRAP+ osteoclasts lining the woven bone along the callus surface, noted by red arrows (B’, B”). (C) The Col1-tk mice also had osteoclasts lining some woven bone, but this was greatly reduced (C’, C”). (D) The ratio of the length of osteoclast surfaces to callus bone surface was significantly lower in Col1-tk compared to WT mice (***p < 0.001). (E) Endomucin staining (brown) of the vessels (orange arrowheads) was prevalent throughout the callus of WT mice. (F) Comparatively, the Col1-tk mice had some staining, but overall reduced vasculature. Black scale bars denote 100 μm unless otherwise noted. (G) Quantification of bone and vasculature within four ROI’s (sampling the entire callus) showed significantly reduced bone and vasculature within the Col1-tk mice as compared to WT (**p < 0.01, ***p < 0.001). (H) When this ROI was defined to only woven bone regions within these fields of view, there was still significantly less % vasculature in the Col1-tk mice (**p < 0.01, ***p < 0.001).

### Radiographic Healing was Impaired in Col1-tk Mice 12 Weeks Post Fracture

To evaluate the effects of transient ablation of proliferating osteoblasts on long-term fracture healing, WT and Col1-tk mice were dosed with GCV for 2 or 4 weeks, the drug withdrawn, and the progression of healing followed until sacrifice at 12 weeks (**Figure 6**). WT mice, dosed for either 2 or 4 weeks, displayed a large callus formed around the fracture site at 2 weeks and this callus condensed over time (**Figure 6A, B**). Based on blinded scoring, nearly all WT fractures had fully bridged after 4 weeks of healing (**Figure 6C, D**). In comparison, while the Col1-tk mice also developed a callus, it was greatly reduced in size compared to WT mice and remained this way over 12 weeks (**Figure 6A, B**). Any small mineralized callus present was visible by week 4 and remained consistent in size thereafter. For the Col1-tk mice dosed with GCV for 2 weeks (**Figure 6C**), between 5 and 12 weeks after fracture only 20 – 65% of femurs appeared fully bridged. Similarly in the Col1-tk mice dosed with GCV for 4 weeks (**Figure 6D**), only 10 – 50% of femurs appeared fully bridged 5 to 12 weeks after fracture. The radiographic scores of WT and Col1-tk mice were significantly different from one another at all timepoints beginning at 2 weeks post fracture (2w GCV: χ^2^, p < 0.05; and 4w GCV: χ^2^, p < 0.01).

**Figure 6.**
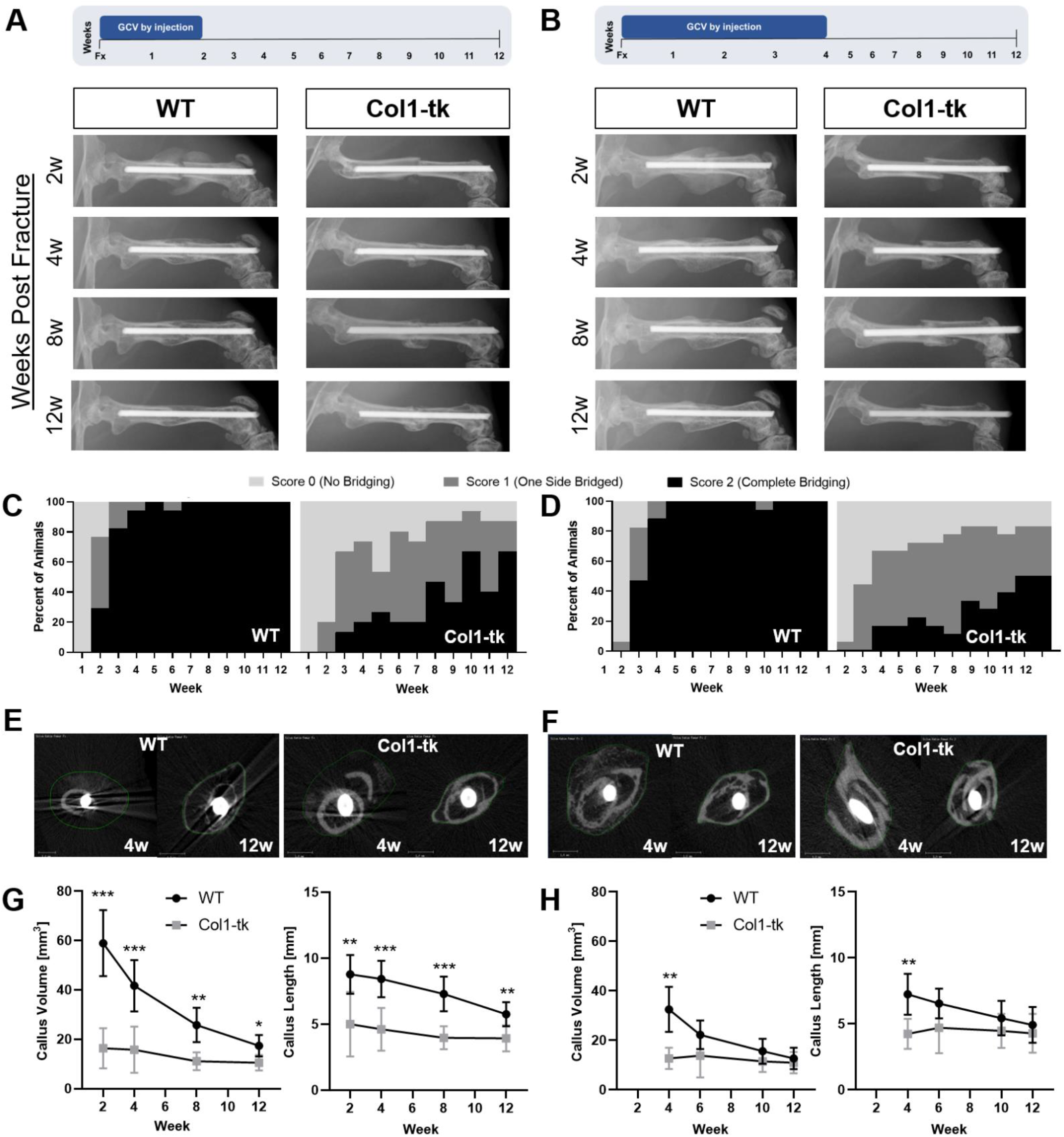
Radiographs were taken weekly following fracture and GCV dosing for (A) 2 or (B) 4 weeks until euthanasia at week 12 (n=12-18/group). Representative X-rays are shown at 2, 4, 8, and 12 weeks for both WT and Col1-tk mice dosed with GCV for either (A) 2 or (B) 4 weeks immediately following fracture. Scoring was completed using a modified Goldberg scale. (C, D) WT mice, regardless of dosing scheme, were fully bridged 4 weeks after fracture. By the end of the study only half of the Col1-tk fractures appeared completely bridged. Scoring of the radiographs demonstrated a significant difference between WT and Col1-tk mice with both 2w (χ2, p < 0.05) and 4w dosing (χ2, p < 0.01). *In vivo* microCT was used to evaluate both callus volume (mm^3^) and length (mm) throughout the 12 weeks of fracture healing with (E, F) representative cross-sections shown from 4 and 12 weeks. (G) Following 2w of GCV dosing, both WT callus volume and length were significantly larger than that of Col1-tk at 2, 4, 8, and 12 weeks. (H) 4w GCV dosing resulted in a significant difference between WT and Col1-tk callus volume and length at the 4 week timepoint. Notably, while WT calluses reduced volume and length over time, there was no significant change in Col1-tk callus size, for either dosing scheme, over time (*p < 0.05, ** p < 0.01, ***p < 0.001).

### In Vivo MicroCT Showed Small Initial Callus Volume that Did Not Change over 12 Weeks in Col1-tk Mice

To further track healing progression through 12 weeks, *in vivo* microCT scans were taken at 2 to 4 week intervals (**Figure 6E, F**). Of the mice dosed with GCV for 2w, WT mice had a significantly larger callus volume and length at all timepoints compared to Col1-tk mice (**Figure 6G**; p < 0.05). In the 4w GCV treated group, WT mice had a larger, longer callus compared to Col1-tk mice only at 4 weeks post fracture (**Figure 6H**; p < 0.01). Callus size in WT mice was greatest at 2 weeks post fracture and then reduced progressively with time, whereas the Col1-tk callus volume and length, for both dosing schemes, did not significantly change over time.

### Ex Vivo MicroCT and Histology Showed Lack of Bridging 12 Weeks Post Fracture in Col1-tk Mice

After 12 weeks of healing, animals were euthanized and both femurs dissected for microCT and biomechanical analysis (**Figure 7**). In WT mice, regardless of dosing scheme, the fracture site was completely bridged by a thin outer cortical shell at the margin of a consolidated callus, and a second inner cortex of similar outer diameter as the original cortex. There were no visible fracture surfaces. Calluses in the WT mice had similar BV as intact bones, but significantly greater TV (**Figure 7B, E**), indicating comparable bone mass distributed over a larger volume. In addition, WT calluses had reduced vBMD and TMD compared to intact femur for both dosing schemes (p < 0.001). By contrast, in Col1-tk mice much of the original cortical bone was still fragmented, with visible fracture surfaces and lack of bridging (**Figure 7B, E**). The callus region in Col1-tk mice had significantly greater BV and TV compared to the intact femur for both dosing schemes (p <0.01), indicating greater bone mass distributed over a larger volume. Consequently, the fractured Col1-tk femurs had similar vBMD as intact bone. TMD was reduced in Col1-tk fractured femurs compared to intact, similar to WT, suggesting callus bone is not fully mineralized in either group. Finally, similar to results at 3 weeks, there were no differences in microCT outcomes in the intact bones of Col1-tk versus WT mice, indicating no effects of targeted cell ablation on intact femora.

**Figure 7.**
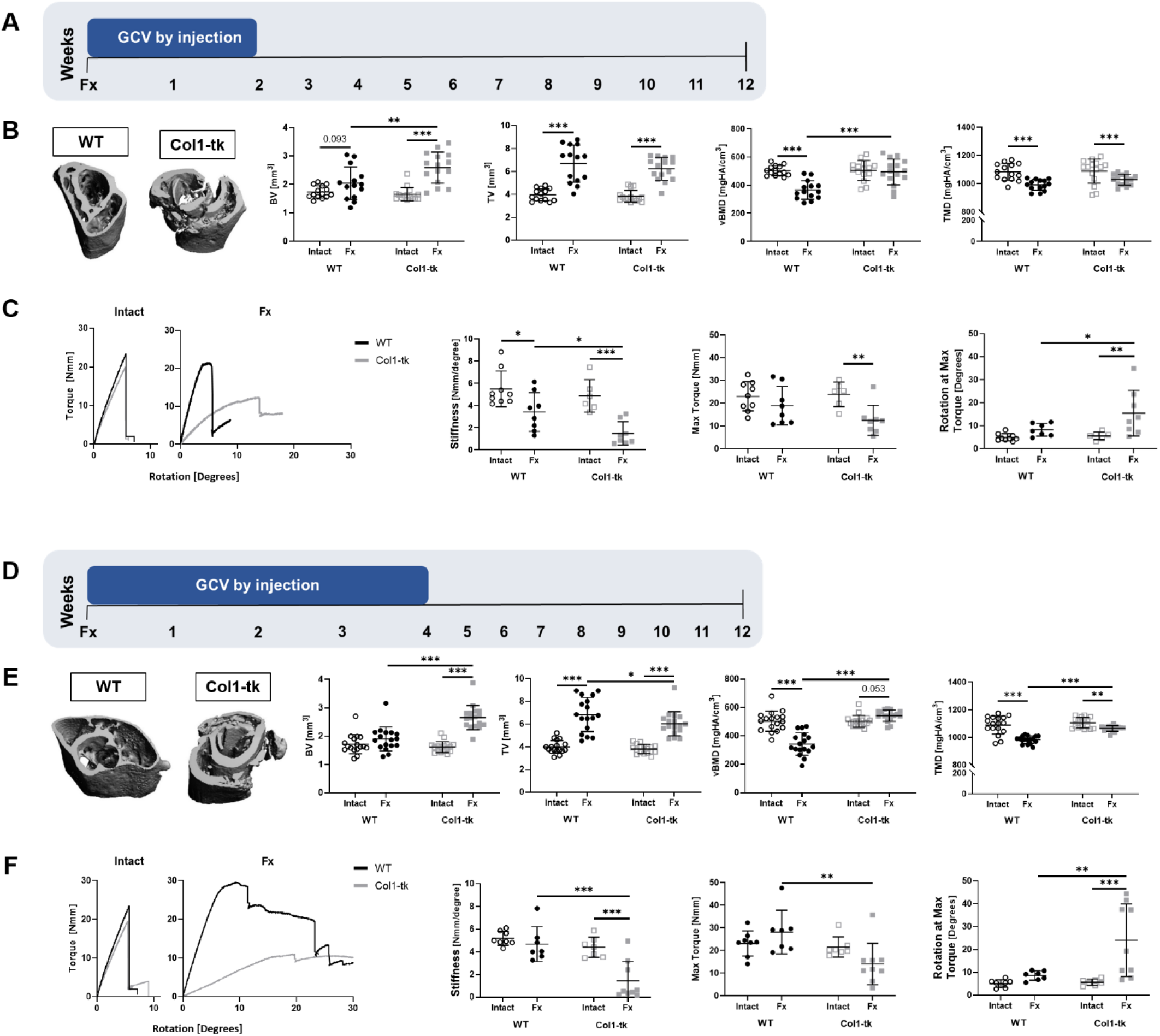
*Ex-vivo* microCT data for WT and Col1-tk intact and fractured (Fx) femurs for both (A) 2 and (D) 4w GCV dosing, analyzed at 12 weeks post fracture (n=15-18/group). (B, E) Representative micro-CT 3D reconstructions displayed a thin, contiguous outer cortex integrated with an inner cortex in WT mice for both dosing schemes. In comparison, the Col1-tk reconstructions show evidence of the original cortical bone with poor consolidation and incomplete bridging at the callus site. (B) For 2w GCV, WT mice had significantly different TV, vBMD, and TMD between intact and Fx femurs. Col1-tk had significantly different BV, TV, and TMD between intact and Fx femurs. (E) Similarly, 4w GCV also resulted in significantly different TV, vBMD, and TMD between the WT intact and Fx femurs. The Col1-tk mice also had significantly different BV, TV, and TMD between the intact and Fx femurs. Finally, the WT and Col1-tk Fx femurs had significantly different BV and vBMD for 2w GCV dosing and BV, TV, vBMD, and TMD for 4w GCV dosing (*p < 0.05, ** p < 0.01, ***p < 0.001). Biomechanical testing was completed to assess fracture stability and strength for (C) 2w and (F) 4w GCV dosed intact and fractures (Fx) femurs, analyzed at 12 weeks post fracture. Representative torque-displacement curves are shown. Regardless of dosing scheme, intact femurs failed before reaching a rotation of 10 degrees. (C) In WT mice, 2w GCV intact and Fx femurs had significantly different stiffness. In Col1-tk, 2w GCV, stiffness, maximum torque, and rotation at maximum torque were all significantly different between intact and Fx femurs. (F) For the Col1-tk, 4w GCV, the stiffness and rotation at maximum torque were significantly different between intact and Fx femurs. Comparisons of Fx femurs between genotypes noted significant differences in stiffness and rotation at maximum torque for both GCV dosing timelines (*p < 0.05, ** p < 0.01, ***p < 0.001). Note that there were no significant differences between WT and Col1-tk intact limbs following either dosing scheme.

### Col1-tk Mice Had Inferior Torsional Properties at 12 Weeks Post Fracture

To assess whether a functional union was achieved, torsional testing was completed on both intact and fractured (“healed”) femurs from WT and Col1-tk mice (**Figure 7C, F**). All intact femurs failed by spiral fracture within 10 degrees of rotation (**Figure 7C, F**), with no significant differences between WT or Col1-tk mice, regardless of GCV duration. In WT mice, the maximum torque and rotation at maximum torque of fractured femurs were not significantly different from intact femurs, indicating return to normal function. The only evidence of incomplete recovery in fractured femurs of WT mice was lower than normal stiffness in the 2w (but not 4w) GCV group. In stark contrast, the fractured femurs in Col1-tk mice had significantly reduced stiffness and increased rotation at maximum torque compared to intact femurs for both dosing groups, with decreased maximum torque in the 2w GCV group (**Figure 7C**). Moreover, comparing fractured femurs between genotypes, Col1-tk femurs had significantly lower stiffness and greater rotation at maximum torque for both dosing timelines, indicating more compliant and less stable behavior, while maximum torque was significantly lower for the 4w GCV group (**Figure 7F**), indicating inferior strength. Thus, WT femurs recover their mechanical properties by 12 weeks post fracture, indicating functional healing, while Col1-tk femurs do not.

### Histology Confirmed Incomplete Healing of Col1-tk at 12 Weeks Post Fracture

Histological sections at the fracture site at 12 weeks were stained with picrosirius red/alcian blue, imaged and analyzed (**Figure 8**). All WT femurs had complete bridging with continuous cortical bone; the original fracture surfaces were not obvious. The thin outer and inner cortices seen on microCT were evident histologically as collagen-rich, aligned bone tissue. In contrast, the Col1-tk femurs all had clearly evident fracture surfaces, with regions that looked like original cortical bone. Col1-tk femurs had some calcified areas with varying degrees of cartilage and fibrous tissue, demonstrating incomplete endochondral bone formation. When quantified, Col1-tk mice had significantly less bone and marrow than the WT mice, as well as significantly more cartilage and fibrous/other tissue, regardless of dosing scheme (**Figure 8C**). When the Col1-tk mice were treated for 4 weeks with GCV, the fracture sites contained significantly more fibrous/other tissue and significantly less bone and marrow than those treated with GCV for only 2 weeks. Calluses were also qualitatively assessed at 12 weeks using H&E, pentachrome, and IHC staining (**Supplementary Figures 4 and 5**). For WT mice dosed with GCV for 2 or 4 weeks, only bone tissue was present (**Supplementary Figures 4B and 5B**), whereas in Col1-tk mice, regardless of dosing scheme, multiple tissue types were present (**Supplementary Figures 4C and 5C**). Additionally, Col1-tk dosed with GCV for 2 weeks displayed areas of positive staining for col2 (within the cartilage region), col3 (fibrous region), TRAP (fibrous/other), and endomucin (all regions). HSV-tk+ cells were found in and around bone, on the margins of cartilage areas and occasionally within fibrous/other tissue areas (**Supplementary Figure 4C**). Comparatively, in Col1-tk dosed with GCV for 4 weeks, HSV-tk+ cells were found sparingly in and around bone (**Supplementary Figure 5C**). There was still evidence of col2 and col3 staining with various tissue types in Col1-tk, but not WT samples.

**Figure 8.**
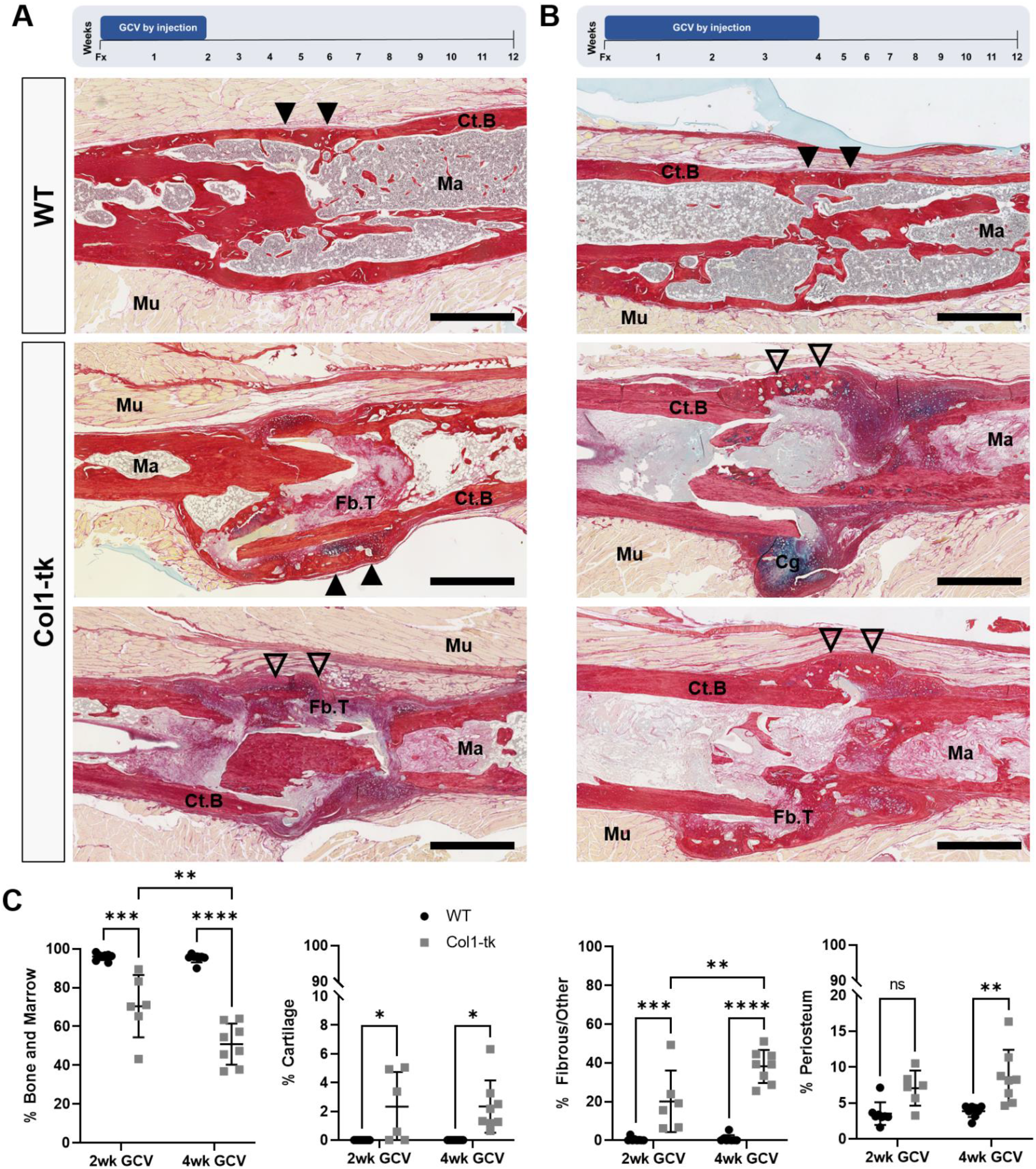
Representative Picrosirius Red/Alcian Blue staining of the WT and Col1-tk fracture site following (A) 2 and (B) 4 weeks of GCV dosing at 12 weeks post fracture (n=7-8/group). Regardless of GCV dosing timeline (2 or 4 week), WT mice had consistent, complete bridging by 12 weeks as shown by continuous cortical bone (bright red staining; filled arrowheads), with little evidence of the original fracture site. The Col1-tk femur calluses had variable healing at 12 weeks (shown by two examples for each dosing group). The callus had some cortical bridging (filled arrowheads) in addition to pockets of woven bone at a still visible fracture site (open arrowheads). The Col1-tk fracture sites had some persistent cartilage and fibrous tissue. Statistical analysis demonstrated significantly less Bone and Marrow, significantly more Cg, and significantly more Fb.T/Other in the Col1-tk mice as compared to WT, regardless of dosing scheme. Additionally, 4w GCV dosing had significantly less Bone and Marrow and significantly more Fb.T/Other as compared to the 2w GCV dosing scheme (*p < 0.05, ** p < 0.01, ***p < 0.001). Abbreviations: Cortical Bone = Ct.B,; Cartilage = Cg; Fibrous Tissue = Fb.T.; Marrow = Ma; Muscle = Mu. Black scale bars denote 1 mm.

When stained for TRAP+ osteoclasts, both the WT and Col1-tk callus periphery had some positive staining at 12 weeks (**Supplementary Figure 6A, A’, A”, B, B’, B”**). The periphery of the callus of WT mice was composed of bone (100% of callus length), while the callus in Col1-tk was a mix of bone and fibrous tissue (73% and 27%, respectively; **Supplementary Figure 6C**). Quantitative analysis of osteoclast length per callus length revealed fewer osteoclasts covering the WT callus than at 3 weeks (**Figure 5**), although still significantly more than Col1-tk (p < 0.05; **Supplementary Figure 6D**). Endomucin staining revealed almost no vessels at the original site of fracture in WT mice (**Supplementary Figure 6E, F**) and minimal vasculature in Col1-tk mice (**Supplementary Figure 6G, H**).

## Discussion

We hypothesized that proliferation of periosteal osteoblast-lineage cells is required for fracture healing. Through a comprehensive assessment of healing in Col1-tk mice, we provide evidence that ablation of proliferating osteoblast-lineage cells during the initial weeks of fracture healing blunts callus formation, leading to impaired radiographic and biomechanical healing. This study therefore establishes the Col1-tk mouse as a new model of atrophic nonunion.

Periosteal cell proliferation is well documented during fracture healing,^(11,14,15,36)^ and expansion of the mesenchymal progenitor cell population from the periosteum appears to account for the formation of cartilage and bone callus.^(21,37)^ However, a requirement for osteoprogenitor cell proliferation has not been directly tested. Previous studies have identified a peak in the proliferative response following fracture that extends from 2 to 14 days, suggesting a critical proliferative window for proper healing.^(14–16)^ But the molecular identity of the proliferating cell populations that contribute to the periosteal callus are not known. Recent work demonstrates that proliferation of 3.6Col1a1 expressing cells contributes to periosteal bone formation after non-injurious mechanical loading.^(11,20,38)^ Herein, using the Col1-tk mouse, we have shown that proliferation of 3.6Col1a1 expressing cells is essential for fracture healing. By targeting the proliferation of these cells and ablating them through exposure to GCV, cell proliferation and fracture callus formation were markedly reduced at early timepoints. Upon withdrawal of the drug after 2 or 4 weeks, the Col1-tk mice were unable to heal 12 weeks post fracture, analogous to what is seen clinically in atrophic nonunion patients. Note that proliferation was still observed in Col1-tk mice, and there was a small mineralized callus evident radiographically and histologically. This may be explained by less than 100% efficiency of the tk-GCV strategy to ablate proliferating 3.6Col1a1 expressing cells, or may reflect contribution of a non-3.6Col1a1 osteoprogenitor cell population. It is also possible that these callus fragments derive from endochondral ossification whereby post-mitotic chondrocytes transform directly to osteoblasts.^(36)^ Nevertheless, the striking reduction in callus formation, impaired callus mechanical strength, and overall inability to recover from the dosing-induced nonunion demonstrates the critical role of 3.6Col1a1 cell proliferation in fracture healing and further supports their essential function in bone formation.^(18,20)^

The 3.6Col1a1 promoter has reported expression in a range of cell types, including osteoblasts, osteocytes, periosteal fibroblasts, tendon cells, and cells of the osteoclast lineage,^(18–20,22)^ raising the question of which cell types were targeted in our study. In fracture callus of 3.6Col1-tk mice, we observed expression of HSV-tk in osteoblasts and new osteocytes, as well as peripheral chondrocytes in cartilage regions and some cells in fibrous tissue and in expanded periosteum. We did not see HSV-tk in chondrocytes in the interior of cartilage regions, or in blood vessel (endomucin+) or osteoclasts (TRAP+). The pattern of HSV-tk expression in callus, and the potent inhibition of bone formation in Col1-tk mice treated with GCV, indicates that proliferating osteoblast lineage cells were targeted for ablation. These results are supported by the limb bud assay, which showed strongly impaired mineralization of Col1-tk cells in presence of GCV (day 16), and are consistent with our recent report of blunted bone formation induced by non-injurious mechanical loading in Col1-tk mice.^(20)^ In contrast, our results indicate that cartilage formation was only modestly affected in Col1-tk mice, evident by grossly normal chondrogenesis in the limb assay (day 6) and by presence of cartilage regions in fracture callus. This is consistent with HSV-tk expression being localized to peripheral (likely hypertrophic) chondrocytes near cartilage-bone transitions in the callus. Therefore, we attribute impaired fracture healing in the Col1-tk mice primarily to ablation of proliferation cells of the osteoblast lineage.

Despite recognition that cell proliferation is a hallmark of fracture healing, the critical window for its contribution to successful healing has not been identified. In rodents, proliferation peaks during the first 2 weeks after fracture, and complete bone bridging and recovery of mechanical properties takes approximately 4 weeks.^(14–16,23)^ We therefore selected 2 and 4 weeks for the durations of GCV dosing. We anticipated that blocking osteoblastic cell proliferation during the entire 4-week healing time course would lead to a permanent nonunion. In contrast, we were unsure whether withdrawal of GCV after 2 weeks might lead to a “rebound” of cell proliferation and eventual (delayed) union. Our results show that the two dosing schemes led to similar outcomes; both prevented healing over the 12 week study duration and led to nonunion. Thus, in mice, proliferation during the first 2 weeks of healing is essential to a full recovery, and a 2-week course of GCV in Col1-tk mice is sufficient to induce nonunion. This finding suggests two possibilities: the local signals that drive cell proliferation are transient, or the proliferative potential of osteoprogenitors is exhausted after 2 weeks of GCV treatment.

One implication of this study is that we have established conditions for an atrophic nonunion without additional local injury (beyond the initial fracture). Clinically, atrophic nonunions are characterized by sparse or absent radiographic callus with primarily fibrous tissue at the fracture site, although cartilage and bone can also be present.^(4,10,39–41)^ Available evidence also supports that atrophic callus tissue is not uniformly avascular.^(10,40)^ The radiographic and histological findings in Col1-tk mice have strong concordance with clinical observations, including minimal radiographic callus, heterogeneous callus tissue with majority fibrous tissue but also some cartilage and bone, and both avascular and vascularized regions. There are many described models of atrophic nonunion in large animals and rodents created by mechanical or thermal disruption of periosteum and bone at the fracture site.^(6,8,26,42–50)^ For example, Garcia et al. ^(26)^ created approximately 2 mm segmental defects with periosteal stripping in mice femora, and observed atrophic nounion after 15 weeks. A recent study by Wang et al. ^(51)^ demonstrated a nonsurgical atrophic nonunion model using radiation to induce periosteal damage. As fracture is a late side effect of radiation therapy, their model provides an appropriate model of the delayed healing response in that clinical setting. However, while these approaches result in atrophic nonunion, such invasive methods are not always representative of clinical nonunions where osseous regeneration has been arrested by a disturbance of metabolic pathways.^(1,9,10)^ Thus, a need has remained for the development of less invasive, pre-clinical atrophic nonunion models to study underlying biological processes and to test therapeutic interventions. We propose the Col1-tk mouse as such a model, with a “failure of biology” that mimics many features of clinical atrophic nonunions.

There are several limitations to the use of the Col1-tk model as presented, as well as need for future studies. First, as noted above, the use of the 3.6Col1a1 promoter to drive HSV-tk targets multiple cell types, and has limitations for targeting well-defined osteoblast lineage cells. To study the contribution of proliferating cells with greater precision will require new tools in which HSV-tk can be driven by more well-defined promoters. Second, due to the ability of the 2 week GCV dosing scheme to prevent recovery from the fracture and lead to development of a nonunion, it is of interest to explore shorter windows of proliferative ablation. A shorter dosing scheme could be used to further examine the potential for rescue of callus formation. Additionally, the dosing start time could be delayed to evaluate the point at which fracture healing progression has gained enough momentum to not be affected by this proliferation ablation. Third, this fracture model used an intramedullary pin for fracture stabilization. Other options include intramedullary locking nail or compression screw, external fixator, a pin-clip device, and locking plates.^(52)^ It is important to consider potential stress shielding with various models as it can cause asymmetric callus formation or even prevent a periosteal response.^(53)^ Fourth, it would be of interest to vary the fracture model by both type and location. While the femur is accepted as appropriate for studying fracture healing, other bones such as the tibia, ulna, rib, radius, and mandible could be useful alternatives.^(52)^ In addition, the full fracture model from this study heals by both intramembranous and endochondral healing.^(28,29)^ A stress fracture model could also be used to focus primarily on intramembranous ossification.^(54,55)^

In conclusion, we have identified a critical role for proliferation of the 3.6Col1a1 cell population in the first 2 weeks of fracture healing, and developed a novel murine model of atrophic nonunion. We utilized a Col1-tk mouse model that, when dosed with GCV for 2 or 4 weeks, effectively blunted callus formation, leading to radiographic and biomechanical nonunion 12 weeks after fracture. The importance of our study is supported by the clinical prevalence of atrophic nonunion, and the expressed need for a relevant model to further study the biology of nonunion. This model can be used in future studies to test new intervention techniques to rescue atrophic nonunion.

## Acknowledgements

This work was supported by funding from NIAMS (R01 AR050211, P30 AR057235, R21 AR076636-01, T32 AR060719, and F32 AR076191-01). The authors would like to thank the Washington University in St. Louis Musculoskeletal Research Center (MRC) Cores and staff for assistance. Specifically, thanks to Yung Kim for all X-ray and microCT (Scanco) acquisition assistance. Also thanks to both Crystal Idleburg and Samantha Coleman for histological processing and sectioning of all specimens. Thanks also to Dennis Oakley of the Washington University in St. Louis Center for Cellular Imaging (WUCCI) Core and Heather Zannit for training and frozen section imaging assistance. Paraffin histological images were taken with the Nanozoomer at Alafi Neuroimaging Core (S10 RR027552). Finally, thanks to Nicole Migotsky for torsion testing and LabVIEW software instruction, as well as Hongjun Zheng and Lisa Lawson for limb bud assay assistance. 3.6Col1a1-tk mice were kindly provided by the labs of Drs. Robert Jilka and Charles O’Brien (University of Arkansas for Medical Sciences, Little Rock, AR, USA).

## Authors’ Roles

Study design: KRH, JAM, AM, MJS. Study conduct: KRH, DAWS, SY, JAM, AH, HZ, DS. Data collection: KRH, DAWS, SY, JAM, EGB, HZ, DS. Data analysis: KRH, DAWS, SY, JAM, AH, EGB. Data interpretation: KRH, JAM, AM, ANM, MJS. Drafting manuscript: KRH. Revising manuscript: KRH, JAM, MJS. Approving final version of manuscript: KRH, DAWS, SY, JAM, AH, EGB, HZ, DS, AM, ANM, MJS. KRH takes responsibility for integrity of data analysis.

