## Supplemental Figures 1-6, Table 1 for "Ablation of Proliferating Osteoblast Lineage Cells After Fracture Leads to Atrophic Nonunion in a Mouse Model"

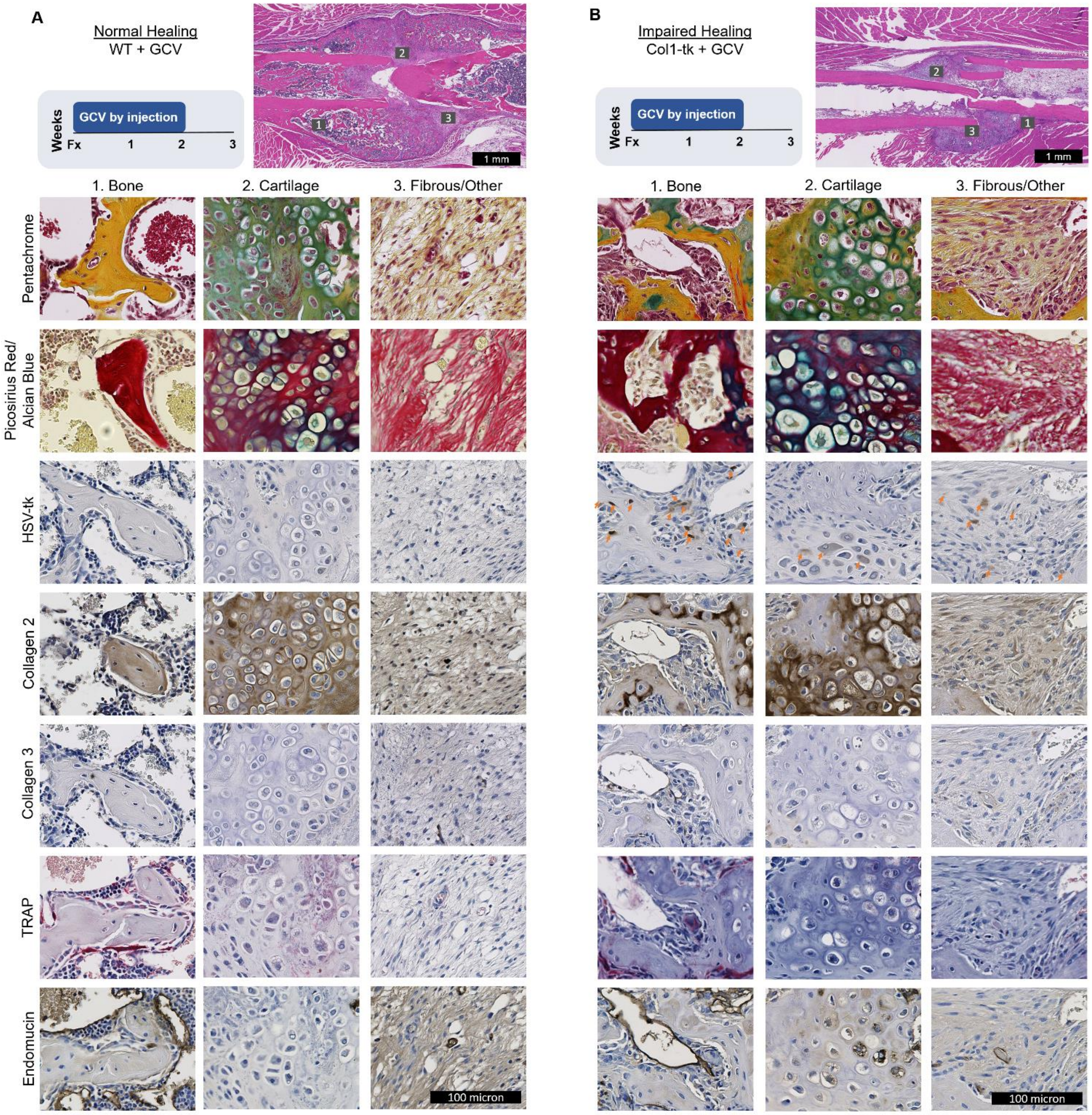


**Supplementary Figure 1.** Histological characterization of callus 3 weeks after fracture for (A) WT and (B) Col1-tk samples. All mice were treated with GCV for 2 weeks. H&E stained callus sections were used to find areas of 1. Bone, 2. Cartilage, and 3. Fibrous/Other tissues to highlight with 20X stained images from serial sections. Staining was done for pentachrome, picorisius red/alcian blue, HSV-tk, collagen 2, collagen 3, TRAP, and endomucin. All tissue types were present in both WT and Col-TK samples. The only staining that differed between WT and Col-TK samples was the HSV-tk. As expected the WT samples had no HSV-TK staining as the gene was not present within the mouse. In the Col-TK sample, similar to Figure 2, the HSV-tk positive cells were found within and around the bone tissue, on the mineralizing margins of the callus, and within some fibrous tissue areas. Within the fibrous/other tissue area, the HSV-tk+ cells do not appear to be positive for collagen 2, collagen 3, TRAP, or endomucin.

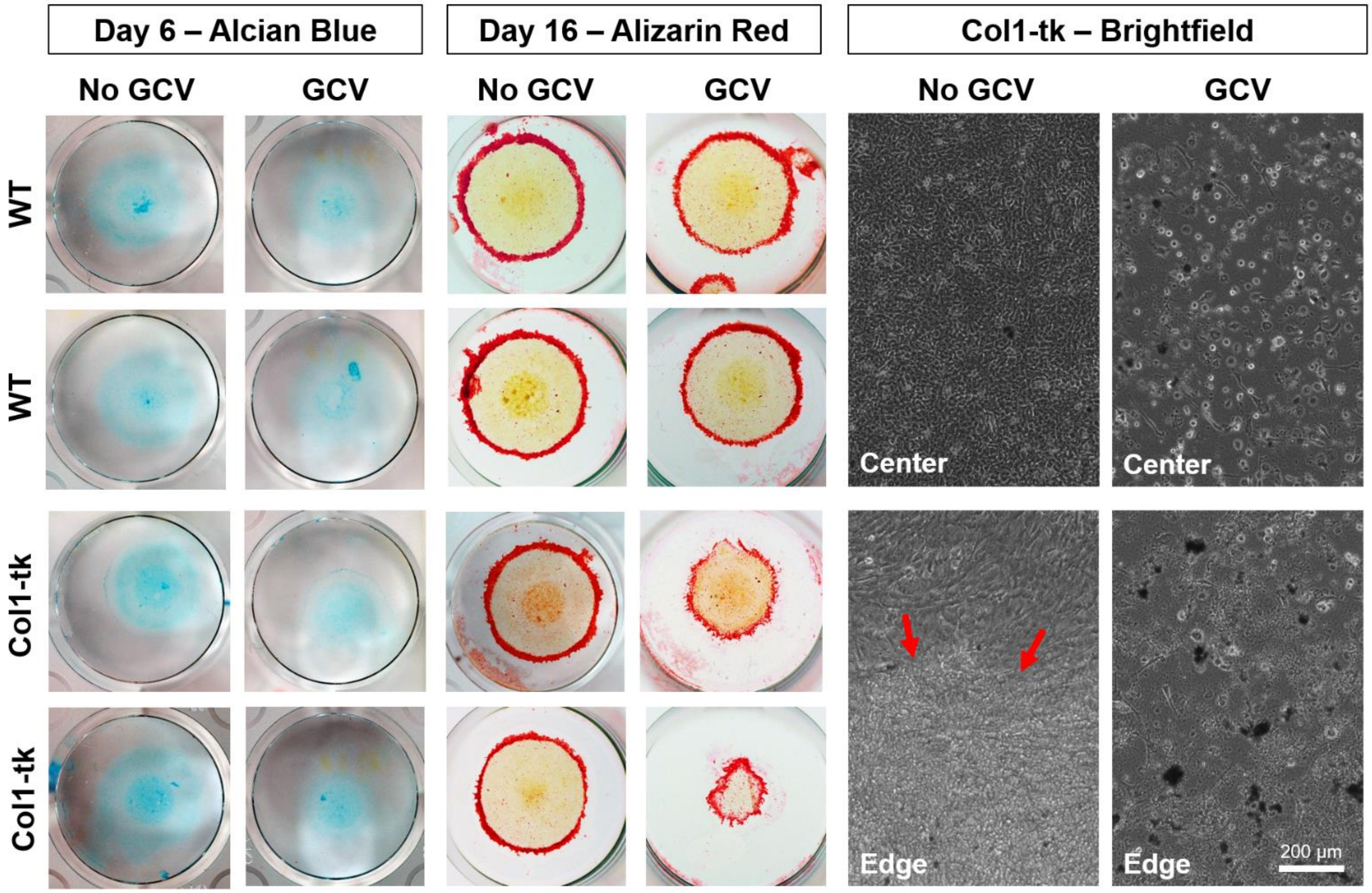


**Supplementary Figure 2.** High-density micromass cultures from murine hind limb buds were utilized to study the effect of GCV/tk on chondrogenesis and osteogenesis. Following isolation, two wells were seeded per sample, with one receiving the addition of GCV to the media. At day 6, there was no difference between WT and Col1-tk cells left untreated; however, when GCV was added to the media, there was a modest effect on chondrogenesis in the cells from Col1-tk mice as shown by slightly reduced staining area. When the cells were given osteogenic media and stained with alizarin red, the Col1-tk cells treated with GCV displayed impaired osteogenesis, as shown by reduced micromass size and overall staining. This is also shown with brightfield images, where Col1-tk cells without GCV cell morphology appeared confluent (center) and had formed mineral clusters at the edge of the well (red arrows), but with the addition of GCV many cells appeared dead throughout (both center and edge) the well.

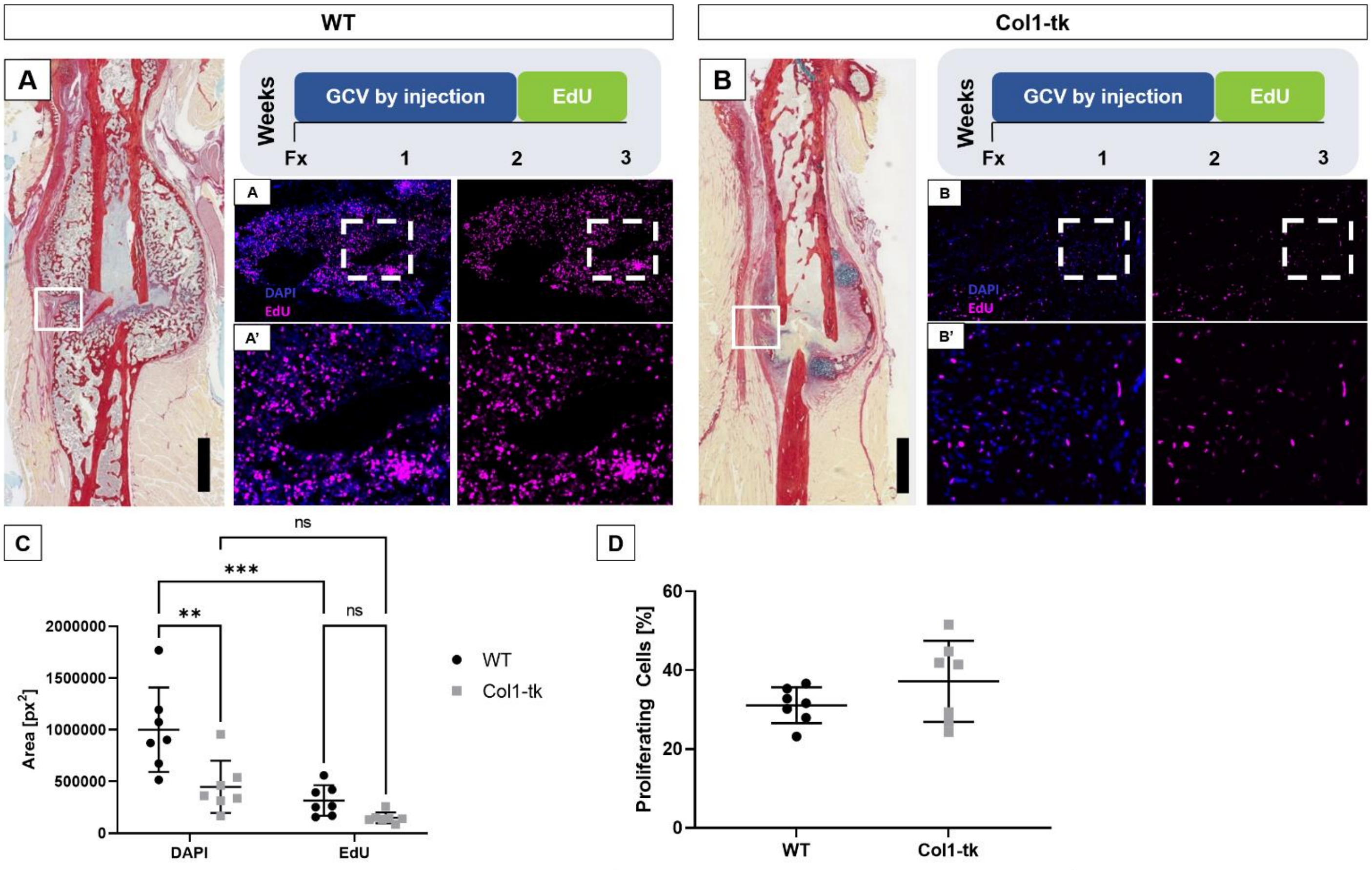


**Supplementary Figure 3.** (A, B) WT and Col1-tk mice were dosed with GCV for 2 weeks, the drug withdrawn, and given EdU for 1 full week post fracture. While it visually appeared that (A, A') WT mice had more EdU+ (pink) staining than the (B, B') Col1-tk mice, quantitative analysis demonstrates that (C) there were no significant differences between the area of EdU+ cells between genotypes. (C) While WT mice had significantly more cells in the fracture callus than the Col1-tk, (D) both had approximately 35% of their total fracture callus cells proliferating. Black scale bars denote 1 mm.

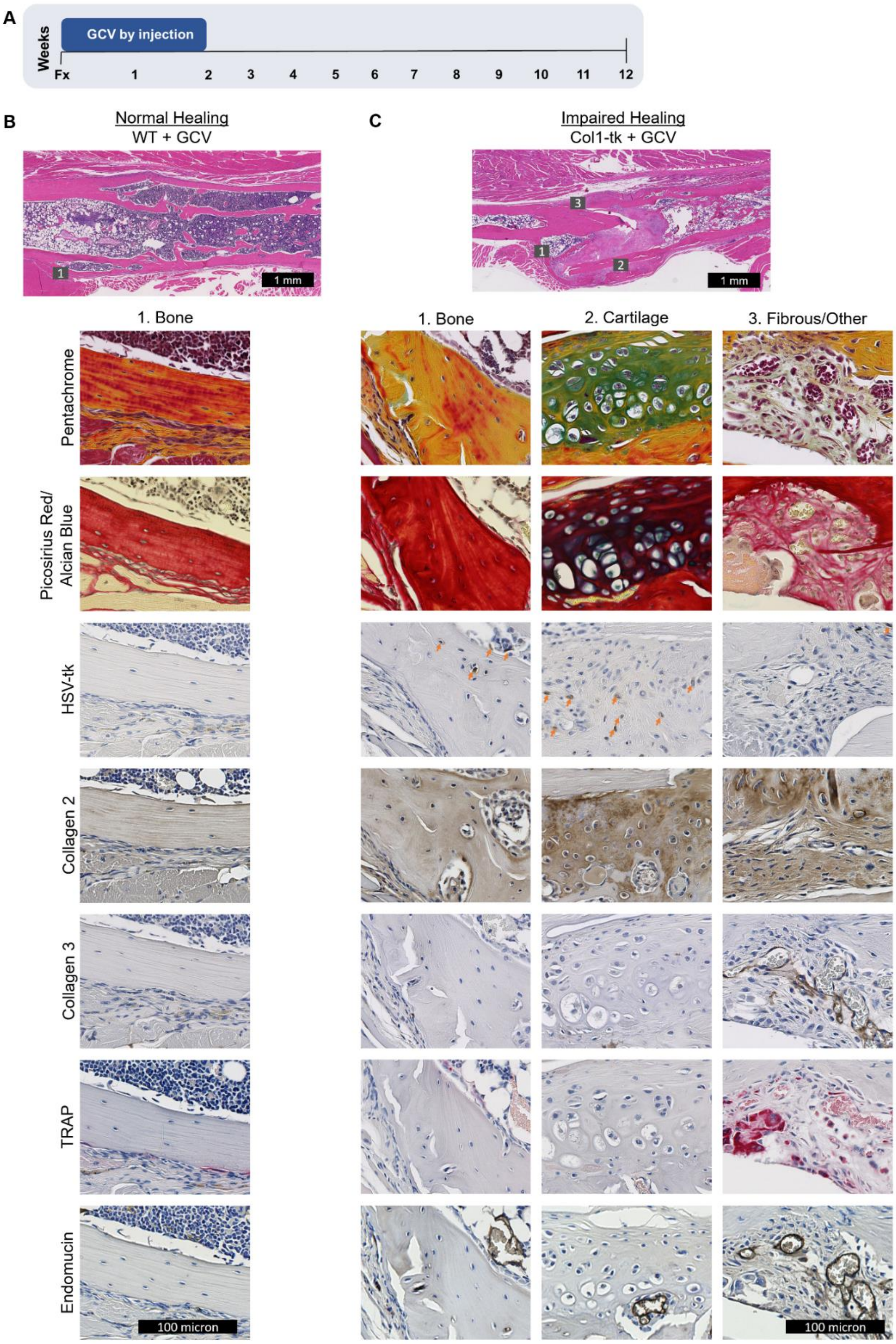


**Supplementary Figure 4.** (A) Histological characterization of callus 12 weeks after fracture for (B) WT and (C) Col1-tk samples. All mice were treated with GCV for 2 weeks. H&E stained callus sections were used to find areas of 1. Bone, 2. Cartilage and 3. Fibrous/Other tissues to highlight with 20X stained images from serial sections. Staining was done for pentachrome, picorisius red/alcian blue, HSV-tk, collagen 2, collagen 3, TRAP, and endomucin. In WT mice only bone tissue was present at the 12 week timepoint, while for Col-TK samples all tissue types were present. WT samples had almost no staining of collagen 2, collagen 3, or endomucin in or around the bone area. There were some TRAP+ cells along the periosteal side of the bone. In the Col-tk samples there were areas of positive staining for collagen 2 (cartilage region), collagen 3 (fibrous region), TRAP (fibrous/other), and endomucin (within all regions). HSV-tk+ cells were found sparingly in and around bone, on the margins of cartilage areas and within fibrous/other tissue areas.

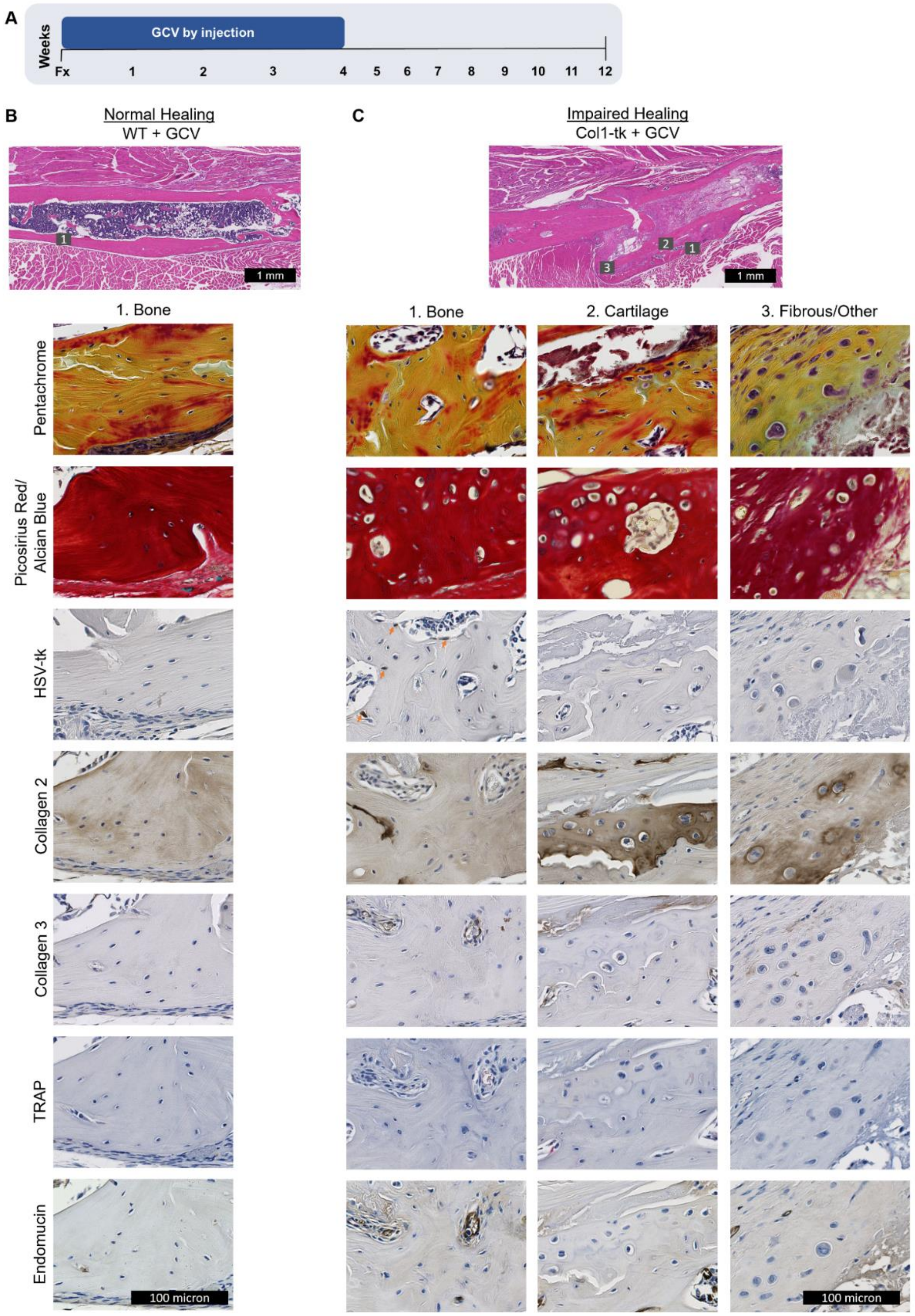


**Supplementary Figure 5.** (A) Histological characterization of callus 12 weeks after fracture for (B) WT and (C) Col1-tk samples. All mice were treated with GCV for 4 weeks. H&E stained callus sections were used to find areas of 1. Bone, 2. Cartilage and 3. Fibrous/Other tissues to highlight with 20X stained images from serial sections. Staining was done for pentachrome, picorisius red/alcian blue, HSV-tk, collagen 2, collagen 3, TRAP, and endomucin. In WT mice only bone tissue was present at the 12 week timepoint, for Col-tk samples all tissue types were present. HSV-tk+ cells were found sparingly in and around bone. There was still evident collagen 2 and collagen 3 staining with various tissue types of Col-tk, but not WT samples.

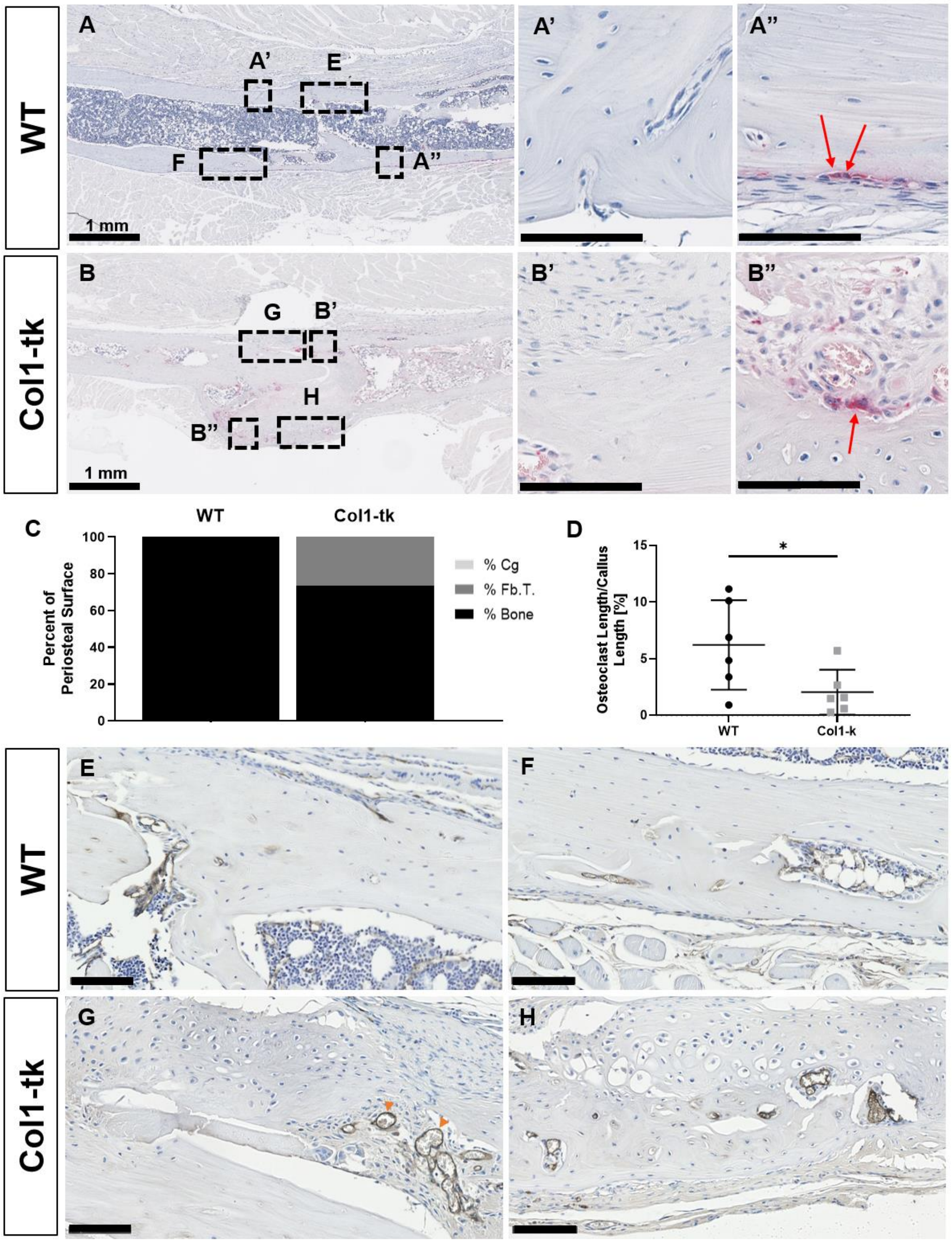


**Supplementary Figure 6.** Osteoclast (TRAP+) activity and the presence of vessels was evaluated in WT and Col1-tk mice at 12 weeks post fracture. (A) WT mice displayed minimal TRAP+ woven bone lining osteoclasts (pink) along the original fracture site, noted by red arrows (A', A"). (B) The Col1-tk mice also had minimal osteoclast activity (B', B"). (C) The callus surface of WT mice was entirely composed of bone and (D) the ratio of the length of osteoclasts to woven bone was significantly higher as compared to Col1-tk mice (*$p < 0.05$). (C) The Col1-tk callus surface was also primarily composed of bone with some fibrous tissue (Fb.T). (E, F) There was very little endomucin staining (brown) of vessels in WT mice. (G, H) While the Col1-tk mice had some endomucin staining, noted by orange arrowheads, overall the presence of vessels was reduced from 3 weeks. Black scale bars denote 100 µm unless otherwise noted.

**Supplementary Table 1.** Cortical morphology and mineral density at the mid-shaft of intact femurs determined by microCT. Data are from mice treated with GCV for 2 weeks and sacrificed 3 weeks after right femur fracture.

| Left Femur (non-fractured; intact) | Total Area ($mm^2$) | | | Bone Area ($mm^2$) | | | Bone Area/Total Area | | | TMD | | |
|---|---|---|---|---|---|---|---|---|---|---|---|---|
| | Mean | SD | P-value | Mean | SD | P-value | Mean | SD | P-value | Mean | SD | P-value |
| WT (n = 12) | 1.73 | 0.164 | NS | 0.869 | 0.109 | NS | 0.500 | 0.022 | NS | 1062 | 38.0 | NS |
| Col1-tk (n = 10) | 1.73 | 0.167 | NS | 0.878 | 0.093 | NS | 0.507 | 0.015 | NS | 1062 | 34.2 | NS |
